# Generation of a new immunodeficient rat model of retinal degeneration with LSL TdTomato reporter and TdTomato-Pcp2 expression

**DOI:** 10.64898/2026.03.04.709611

**Authors:** Magdalene J. Seiler, Helios Nguyen, Devan Endejan, Bin Lin, Guojun Zhao, Lauren Klaskala

**Affiliations:** Department of Physical Medicine & Rehabilitation; Sue & Bill Stem Cell Research Center; University of California, Irvine, U.S.A; Department of Ophthalmology & Visual Sciences, Gavin Herbert Eye Institute, University of California, Irvine, CA, U.S.A; Department of Anatomy & Neurobiology, University of California, Irvine, CA, U.S.A; Brunson Center for Translational Vision Research, University of California, Irvine, CA, U.S.A; Inotiv, Maryland Heights, MO, 63043, USA

**Keywords:** Photoreceptor degeneration, transgenic rat, cre-recombinase, cre-lox cell label, bipolar cells

## Abstract

**Purpose:** To develop an immunodeficient retinal degenerate (RD) rat model with fluorescent label for studying retinal degeneration and transplant-host connectivity.

**Methods:** Gene constructs for *CAG-LSL-TdTomato* and *Pcp2-Cre* were developed and injected into rat embryos at Envigo. The LSL TdTomato reporter strain, created on immunodeficient *RhoS334ter-3* rats (RRRC#539), was bred to homozygosity at UCI (strain *SD-Foxn1rnuTg((Rho-S334X)3,CAG-TdTomato)1010Mjsuc*, “RNT”). The second gene construct *Pcp2-Cre* was injected into Long-Evans (LE) rat embryos, resulting in two *Pcp2-cre* founders (strain *PCP2 Cre-1105 RKI,* “Pcp2”), with targeted and targeted/random insertion. Founders were mated with an LE male and a *foxn1+/-* NIH nude female. F1 offspring was bred to homozygosity and immunodeficiency. Homozygous rats of both strains were crossbred to generate *TdTomato-Pcp2 RD* (“RTP”) rats. Retinas were processed for immunohistochemistry for various retinal markers. GFP-expressing rat retinas were transplanted to 6 week old “RTP” rats and analyzed 37 and 77 days post-surgery (pilot experiments).

**Results:** TdTomato-Pcp2 RD rats exhibit RD similar to the original *Rho S334ter-3* rat strain, with < 1 row photoreceptors remaining at 1 month. Retinas with targeted *Pcp2* insertion showed TdTomato in retinal interneurons and cones. Retinas with random *Pcp2* insertion exhibited additional TdTomato in RPE, glial, and endothelial cells. *Pcp2-TdTomato* expression was useful to define transplant-host boundaries.

**Conclusions:** We created an unique RD rat model for studying retinal transplant connectivity. The RD *LSL-TdTomato* reporter rat can also be used to generate RD rats with other cell-specific labels using the *cre/lox* system.

**Translational relevance:** This newly created rat model is useful for cell therapy studies.

## Introduction

Retinal degeneration, characterized by the progressive loss of photoreceptors and other retinal neurons, comprises a variety of inherited and acquired disorders, such as retinitis pigmentosa (RP) ^1, 2^ or age-related macular degeneration (AMD) ^3, 4^, leading to irreversible visual impairment and ultimately blindness. These conditions affect millions of people in the US across different ages and disease prognosis stages, yet effective treatments remain limited. Current treatments including nutritional supplements ^3, 5^ to slow down the progression of retinal degeneration, do not aim to replace lost photoreceptors. There are also cell therapies already in clinical trials aiming to slow down photoreceptor loss ^6^. Our research targets advanced retinal degeneration when photoreceptor replacement is necessary.

Animal models have played a major role in elucidating the molecular and cellular mechanisms underlying retinal degeneration and in advancing therapeutic strategies, including gene therapy, pharmacological interventions, and cell replacement (recent review ^7^). While numerous mouse models have been developed to mimic specific genetic mutations and pathological features of human retinal diseases, rat models offer distinct advantages due to their larger ocular anatomy, relatively easier surgical manipulation and longitudinal imaging studies ^8–10^. However, the availability of well-characterized rat models for retinal degeneration remains limited, so further studies and experiments are needed to better fit the research purposes and schemes. Rats have advantages over mice in terms of surgical accessibility (larger eyes) and behavioral testing.

Recent advances in regenerative medicine have opened new avenues for therapeutic intervention, with retinal organoid transplantation emerging as a particularly promising strategy ^11–19^. Retinal organoids are three-dimensional, self-organizing structures that are derived from human pluripotent stem cells capable of recapitulating key features of the human retina, including photoreceptors and retinal interneurons ^20–22^. Transplantation of human retinal organoids into degenerated retinas has shown encouraging results in preclinical models, demonstrating the restoration of visual function to some degree, and offers a renewable source for retinal repair ^14, 16–19, 23^.

In this study, we describe the development and characterization of a novel immunodeficient rat model of retinal degeneration. With the emerging genetic engineering advancement, we employed the Cre-loxP recombination system to create a novel transgenic hybrid rat model containing TdTomato-fluorescent retinal bipolar cells. We created a retinal degenerate strain universally expressing CAG-LSL-TdTomato, and another strain, expressing *Pcp2-cre* on a Long-Evans background that was crossed with nude rats to make it immunodeficient. *Pcp2* was expected to label the rod bipolar cells as has been shown in a mouse model ^24, 25^. After crossing both rat strains, the F1 offspring would be either nude (*foxn1^-/-^*) or non-nude (immunocompetent *foxn1^+/-^*), while everyone will possess the transgenic fluorescent retinal component (*TdT-Pcp2*) and exhibit retinal degeneration. Our findings establish this new model as a valuable tool for advancing the understanding of the retinal connectivity between the transplant and host retinal architecture, which will be valuable for the efficiency and feasibility of retinal transplantation as a novel therapeutic approach.

## Methods

### Experimental animals (UCI and Envigo)

Animals were treated in accordance with NIH guidelines for the care and use of laboratory animals, the ARVO Statement for the Use of Animals in Ophthalmic and Vision Research, and under a protocol approved by the Institutional Animal Care and Use Committee of UC Irvine. Twenty-four female and 10 males of the immunodeficient *Rho S334ter-3* rat strain *(SD-Foxn1 Tg(S334ter)3Lav)* were shipped to Envigo in March 2022. At Envigo, fertilized embryos were injected with gene constructs at the 1-cell stage and implanted into pseudo-pregnant females. After identifying a transgenic founder, the F1 generation of the *TdTomato-1010* rats was received in April 2023. *Pcp2-Cre* rats were generated on a Long-Evans (LE) background after embryo injections into immunodeficient *RhoS334ter-3* rats failed. The F1 generation of the *Pcp2-Cre 1105* rats was received in March 2024 (bred with LE and NIH nude rats). Both rat strains were bred to homozygosity at UCI. Homozygous rats of both strains were bred to produce TdTomato expressing “RTP” rats starting in September 2024. Some “RTP” rats were processed for immunohistochemistry at different ages, starting at postnatal d19.

Several of these “RTP” rats were transplanted with rat E19 fetal retinal sheets, derived from a GFP-expressing rat donor (RRRC strain #307) crossed with hPAP *(ACI-Tg(R26_hPLAP)* rats.

### DNA donor construction for TdTomato1010 & Pcp2 (Envigo)

#### Use of ZFN to create TdTomato-loxP

To generate the CAG-LSL-TdTomato construct, the TdTomato was combined with the CAG promoter. ZFN engineering was used to create the rROSA-TdTomato-1042 model, with donor DNA (CAG promoter, LSL-TdTomato reporter). **Figure 1A** shows a diagram for validating transgene constructs. The Zinc finger nuclease (ZFN) technology was used to generate the TdTomato1010 construct. ZFN bound to the region of interest on the DNA sequence and introduced DNA double strand breaks (DSBs). These DSBs underwent natural cellular reparation, via either error-prone nonhomologous end joining (NHEJ) or high-fidelity homologous recombination (HR) with the provided donor DNA. Each ZFN pair was validated by transfection into cultured rat C6 cells, and cutting efficiency assessed by the Surveyor Cel-1 Mutation Detection assay (**Figure 1B**). The two smaller cleavage bands add up to the size of the parental band. The ratio of the intensity of the cleavage bands relative to the parental band is how we assessed cutting efficiency. **Figure 1C** shows the gene construct and the sites for the primers used for genotyping (primers see **Table 1**).

**Figure 1.**
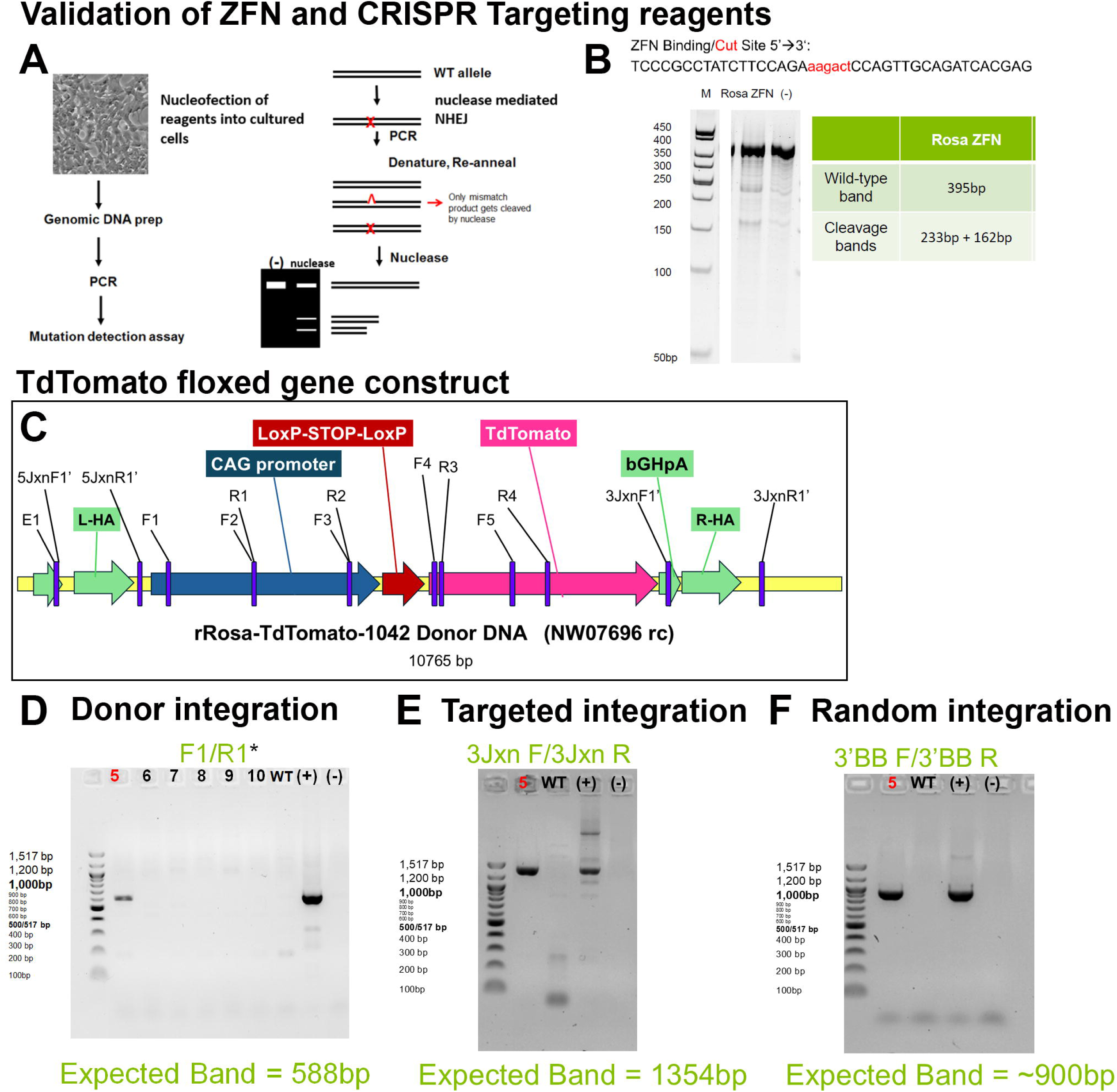
Generation of gene construct for *CAG-LSL-TdTomato* transgene and genotyping of TdTomato 1010 founder. ZFN engineering was used to create the rROSA-TdTomato-1042 model, with donor DNA (CAG promoter, LSL TdTomato reporter). **A)** Schematic diagram of validation strategy for transgene targeting reagents. The Surveyor analysis involves four steps: 1. Preparation of PCR amplicons from the mutant (test) and wild-type (reference) DNA. 2. Hybridization of equal amounts of test and reference DNA by heating and cooling the DNA mixture to induce hetero- and homo-duplex formation. 3. Treatment of the hetero-/homoduplex DNA with SURVEYOR Nuclease. The reference DNA alone serves as background control. 4. Analysis of the DNA fragments by gel electrophoresis. A mismatch between test and reference DNA will result in the formation of cleavage products by the nuclease, and their size indicates the location of the mismatch. **B)** The ZFN binds and cleaves specific DNA sequences in a given genome, generating DNA double strand breaks (DSBs). These DSBs are repaired via either error-prone nonhomologous end joining (NHEJ), or homologous recombination (HR) when a donor DNA is also provided. The two smaller cleavage bands add up to the size of the parental band. The ratio of the intensity of the cleavage bands relative to the parental band shows cutting efficiency. **C)** Schematic diagram of gene construct with sites of primers indicated. **D-F)** After injection into 156 embryos of RRRC strain #539, 1 donor was identified and backcrossed to the original strain. Examples of genotyping results of founder rat with primer for donor integration **(D)**, F1/R1 primer (similar results with other donor integration primers F2/R2, F3/R3, F4/R4), targeted integration **(E)**, 3JxnF1’/3JxnR1’ primer (similar results with 5JxnF1’/5JxnR1’ primer), and random integration (**F)** , primer 3BB F1’/3BB R1’ (similar results with 5BB F1’/5BB R1’).

**Table 1:**
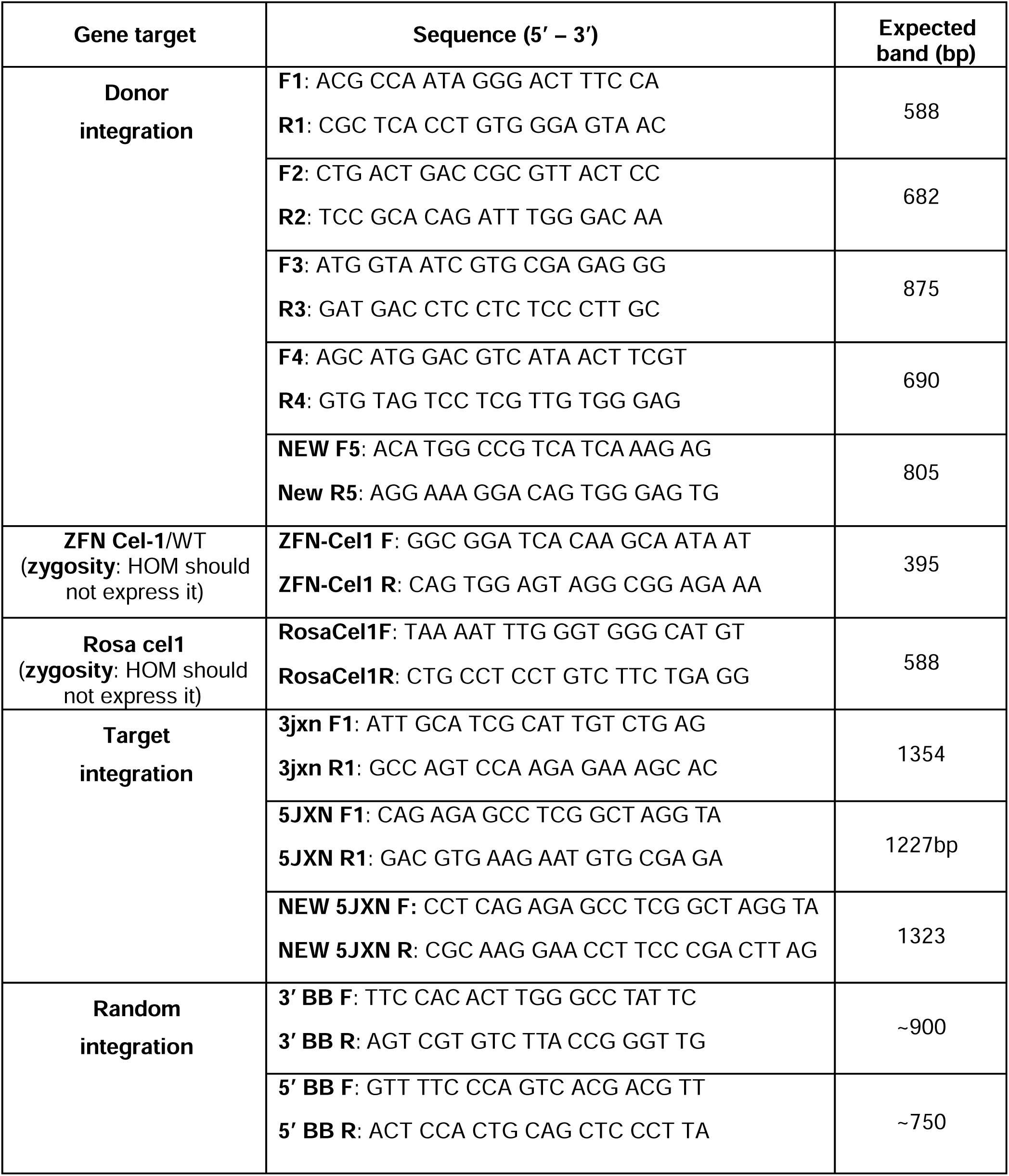
CAG-LSL-TdTomato 1010 primers

The validated gene constructs were then injected into 156 pronuclear embryos of RRRC strain #539 ^10^. One donor was identified and backcrossed to the original strain (genotyping results in **Figure 1D-F**).

*TdTomato1010 Primers* (see **Table 1**): Initial pairings of F1/R1, F2/R2, F3/R3, F4/R4, and F5/R5 primers were used to identify donor integration; 5jxn F/5jxn R to identify targeted integration on the 5’ side; 3jxn F/3jxn R to identify targeted integration on the 3’ side. Potential founders were screened for random integration using the primers 3BB F1’/3BB R1’ and 5BB F1’/5BB R1’.

### CRISPR-Cas9 for Pcp2-Cre

The design of the gRNA is shown in **Figure 2A-C**. gRNAs, donor material, Cas9 mRNA, injection buffer, and Cas9 protein (-20°C) were thawed on ice. gRNAs were lightly vortexed and centrifuged for two minutes at 15,000 x g at 4^°^C. In a low-binding 0.5-ml microfuge tube, appropriate amount of gRNAs, Cas9 protein, and Cas9 mRNA were combined, allowing gRNA and Cas9 protein to complex at room temperature for 15 minutes. A cocktail of DNA donors, Cas9 mRNA, and injection buffer was centrifuged for 10 minutes at 4°C at 14,000 RPM. Roughly 20 µl was removed from the top of the mixture without disturbing the bottom and placed in a new tube without creating bubbles. This 20µl aliquot was used for pronuclear embryo microinjection.

**Figure 2:**
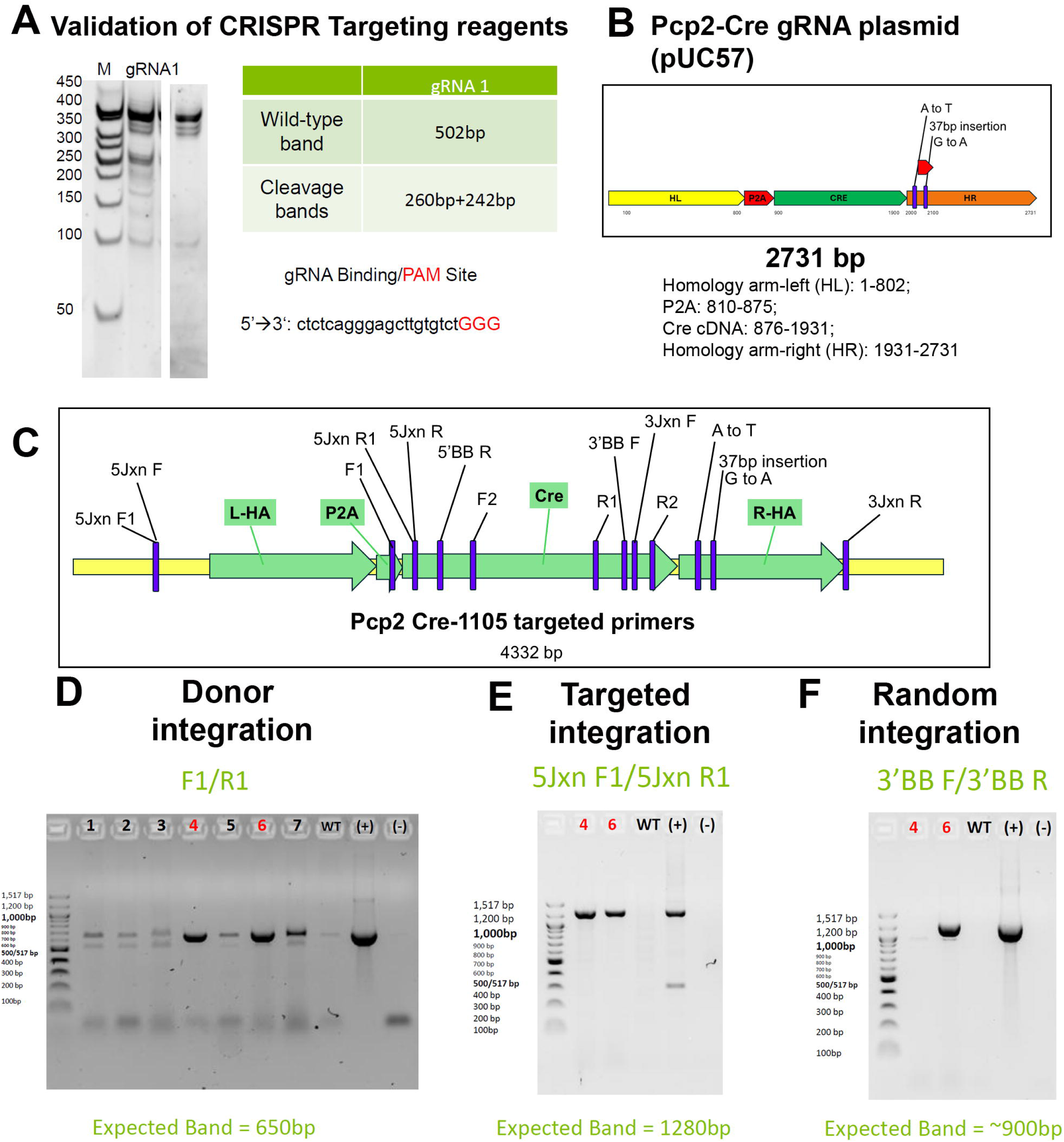
Generation of Pcp2-Cre gene construct and genotyping of Pcp2-Cre **founders**. CRISPR-Cas9 engineering was to create the *Pcp2 Cre1105* donor DNA. **A)** The CRISPR/Cas9-gRNA complex binds and cleaves specific DNA sequences in a given genome, generating double strand breaks (DSBs). The cell then repairs DSBs via either error-prone nonhomologous end joining (NHEJ), or high-fidelity homologous recombination (HR) when a donor is also provided. We validated each small guide RNA by transfection into cultured cells, and assessed cutting efficiency by the Surveyor Cel-1 Mutation Detection assay. The two smaller cleavage bands add up to the size of the parental band. The ratio of the intensity of the cleavage bands relative to the parental band is how we assess cutting efficiency. **B)** Guide RNA plasmid. Homology arm-left(HL): 1-802; P2A: 810-875; Cre cDNA: 876-1931; Homology arm-right (HR): 1931-2731. **C)**. Diagram for Pcp2 Cre-1105 targeted primers (see also Table 2). **D-F)** Genotyping results for the 2 *Pcp2-Cre* founders (after unsuccessful injection into 538 embryos of the RRRC strain #539, two founders were identified after injection of 291 Long-Evans embryos). **D)** Donor integration identified with F1/R1 primer (2 out of 7 animals, indicated in red). **E)** Targeted integration shown by 5Jxn F1/5Jxn1 R1 primers. **F)** One of the founders, animal #6, also exhibited random integration of *Pcp2-Cre* as shown with 3’BB F/3’ BB R primer.

**Table 2:**
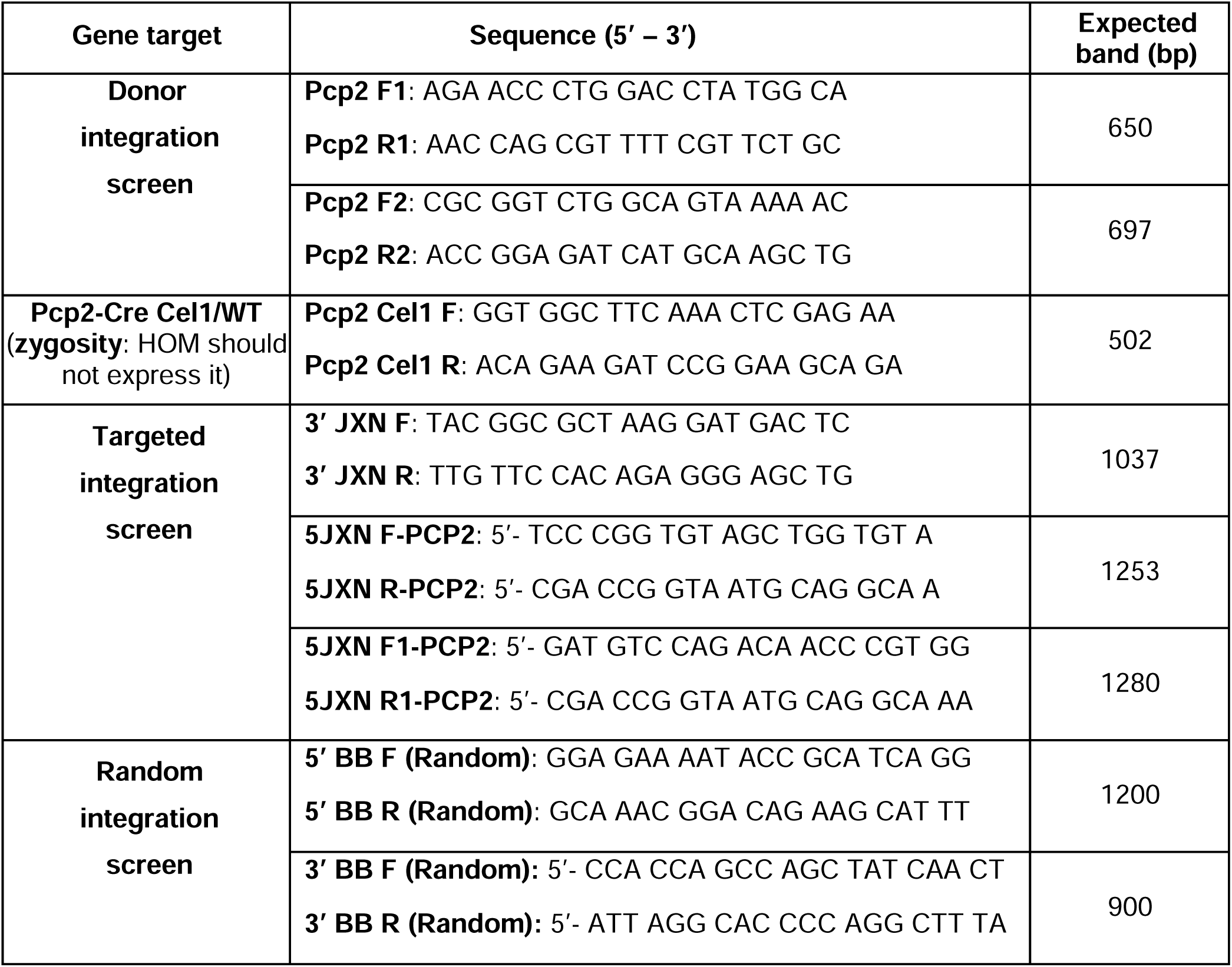
Pcp2-Cre Primers

### Validation of the targeting reagents

Using primers flaking the target site, both the modified and wild-type (WT) sequences were subjected to polymerase chain reaction (PCR) for amplification. The cycle settings allowed the molecules to form both homo and heteroduplexes. Cel-1 endonuclease was utilized to detect heteroduplex and cleaved the mismatched sequences. The digested products were resolved by gel electrophoresis (see **Figure 1** for TdTomato 1010, and **Figure 2** for Pcp2-Cre). Primer sequences are listed in **Table 1** (TdTomato 1010) and **Table 2** (Pcp2-Cre-1105).

### Genotyping

#### DNA extraction

Tail tissues were collected at the same time as tattooing during neonatal period. Tissues were preserved at -80°C until ready to be extracted. DNA extraction method followed the protocol provided by the GenCatch™ Blood & Tissue Genomic Mini Prep Kit (Epoch Life Science).

Tissues were incubated with proteinase K in lysis buffer at 60°C for about 3 hours. Nucleic acid contents were precipitated by absolute ethanol at room temperature. The mixture was then washed with washing buffer and centrifuged at 8000 RPM for 2 minute to remove the organic contents and ethanol residues. Preheated Tris-EDTA (TE) buffer (pH 8.0) was used to elute the DNA content. The eluted DNA was stored at -20°C until ready to be amplified.

### PCR and gel electrophoresis

DNA vials were thawed and kept on ice. Measurements of nucleic acid and proteins concentration was performed by Thermo Scientific NanoDrop™ 2000/2000c Spectrophotometer using a volume of 1uL for each sample. Each Polymerase Chain Reaction (PCR) reaction cocktail contained 12.5ul of the 1X repliQa® HiFi ToughMix® (Quantabio) master mix (MgCl_2_, dNTPs, proprietarily formulated HiFi polymerase, hot start antibodies and ToughMix chemistry), 1ul of 400nM each of Forward and Reverse primers, 50ng of DNA templates, and corresponding RNAse-free water that added up to a total of 25ul per vial.

PCR reactions were run on Eppendorf Mastercycler® (Eppendorf) following these thermal conditions for 35 cycles: initial denaturation at 95°C for 2 min, denaturation 95°C for 45s, annealing at 60°C for 1 min, extension at 72°C for 2 min, final extension at 72°C for 5 min, and holding at 4°C indefinitely. Samples were resolved on 2% agarose gel (TopVision Agarose, Thermo Scientific™) and 1X GelRed® Nucleic Acid Stain (Biotium). Gel was visualized under the UV light using the ChemiDoc XRS+ Imager (Biorad).

### Determination of zygosity

**Table 3** shows the strategy to determine zygosity of rats (this applies both to CAG-LSL-TdTomato and Pcp2-Cre). Essentially, the Cel1 F/R primer is expressed by WT and heterozygous rats, and donor integration primers are expressed by heterozygous and homozygous transgenic rats.

**Table 3:** Determining zygosity of *TdTomato 1010* or *Pcp2-Cre* rats

| Genotype | Primer |  |
| --- | --- | --- |
|  | ZFN-Cel1 F/R (*TdTomato1010*) or<br>Pcp2-Cre Cel1 F/R (*Pcp2-Cre*) | Donor integration |
| WT | + | – |
| HET | + | + |
| HOM | – | + |

#### Transplantation of rat fetal retinal tissue (Pilot experiments)

Retinal tissues were harvested from rat embryos derived from rats universally expressing EGFP (RRRC strain #307, (F344-Tg(UBC-EGFP)F455Rrrc ^26^ crossed with ACI-Tg(R26_hPLAP, RRRC # 817) at embryonic day 19 (E19). Tissues were dissected into rectangular retinal sheets (1-1.3 x 0.6 mm) and stored overnight at 4°C in HibernateE medium with B-27 supplements (Thermo Fisher Scientific) and brain-derived neurotrophic factor (BDNF)/glial cell-line derived neurotrophic factor (GDNF)-loaded poly(lactic-co-glycolic) acid (PLGA) microspheres).

Recipient rats (age 6 weeks) were anesthetized with ketamine/xylazine cocktail (37.5 mg/kg ketamine and 5 mg/kg xylazine). The pupil of the surgery eye was dilated with tropicamide (1%, Somerset). Using a sapphire microblade (World Precision Instruments), a small incision was made just behind the *pars plana* through the sclera, choroid, and retina, followed by a local retinal detachment. Sheets of retinal tissue were loaded into a custom-made implantation tool (US patent #2 159 218) ^27^ and inserted into the subretinal space. The scleral incision was closed with 10-0 sutures. Immediately afterwards, the fundus was examined by a coverslip on the cornea to locate the transplant placement. The surgery eye was then treated with Neomycin and Polymyxin B Sulfates and Bacitracin Zinc B.N.P triple antibiotic eye ointment (Bausch & Lomb). After injection of Buprenex (0.03 mg/kg) and antisedan (0.25 mg/kg) (MWI Veterinary Supply, Cencora, CA), rats were placed in an incubator and monitored for full recovery before they were returned to the cages.

#### **Optokinetic testing** (OKT, visual acuity)

Rats homozygous for the *CAG-LSL-tdTomato* gene were tested for tracking moving stripes of varying widths in a virtual cylinder (OKT apparatus, Cerebral Mechanics, Alberta, Canada) after at least 1 hour of dark adaptation ^27–29^. Six different frequencies (= alternating stripe widths; 0.05-0.45 cycles/degree) were tested for 1 min. ea. in the clockwise and counterclockwise direction, followed by a blank grey screen, as previously described ^19, 27^. Rats were tested monthly. Videos of the rat’s head following the stripe movements were subsequently analyzed <u>in a masked fashion</u> by 2 independent observers. The best visual acuity of the two same-day tests was used for analysis. If there was a discrepancy between the two observers, videos were re-analyzed by a third observer. All testers and video-watchers were blinded to the experimental group assignment of rats. Statistical analysis and graphing data was performed in Graphpad (version 11.0).

### Tissue processing and immunofluorescence assay

Rats were euthanized according to the AVMA guidelines, either by CO2 or by Ketamine/Xylazine overdose, followed by perfusion fixation with chilled 4% paraformaldehyde (PFA) in 0.1 M sodium phosphate buffer for approximately 10 minutes. Eyes were harvested and soaked in 4% PFA overnight after the cornea was removed to maximize tissue preservation. Tissues were washed the next day with 0.1 M sodium phosphate and further dissected until the eye cups were ready to be infiltrated with 30% sucrose and frozen Tissue-Tek® OCT Compound (Sakura). Tissues were cut along the dorsal-ventral axis using Leica Cryostat, and mounted on Superfrost Plus slides (Fisher Scientific). Slides were incubated in HistoVT One (Nacalai, 1:10 dilution, 30 minutes at 70°C) for antigen retrieval, washed with 1X phosphate buffered saline (PBS), and blocked with 20% donkey serum for at least 30 min at room temperature, followed by the incubation of a cocktail of primary antibodies at 4°C overnight (see **Table 4** for antibodies). After several washes with phosphate-buffered saline (PBS) and a second blocking, the slides were incubated for 30 minutes at room temperature with fluorescence tagged secondary antibodies, Alexa Fluor 488 donkey anti-mouse IgG (H+L) and AF647 donkey anti-rabbit IgG (H+L) (Jackson ImmunoResearch). After further washing with PBS, the slides were incubated with DAPI (4’6’-diamidino-2-phenylinode hydrochloride) for 30 min at room temperature and were coverslipped with Vectashield mounting media (Vector Labs). For Peanut agglutinin (PNA) lectin staining, slides were blocked with carbo-free blocking solution (CFBS; Vector labs, Cat. # CSP-5040-125) containing 3% Triton-X100 (TX), followed by biotinylated PNA (Vector labs, Cat. # B-1075-5) diluted 1:200 in CFBS without TX at 4°C overnight. After 3x PBS washes, slides were then incubated in DyLight 649 Streptavidin (Vector labs, Cat. # SA-5649-1; final conc. 5 ug/ml) in CFBS without TX for 1 hour at room temperature, followed by PBS washes. Slides were then stained with DAPI and coverslipped as described above. Slides were imaged using Leica SP8 confocal microscope (Leica Microsystems).

**Table 4:** Antibodies

| Antibody | Species | Specificity | Dilution | Source; Cat. # | RRID |
| --- | --- | --- | --- | --- | --- |
| Primary antibodies |  |  |  |  |  |
| Calbindin | mouse | Horizontal cells | 1:200 | Sigma Aldrich | AB_476894 |
| CRALBP | mouse | RPE + Müller glia (MG) | 1:1000 | Abcam | AB_2269474 |
| GFAP | goat | Astrocytes + MG (reactive) | 1:500 | Sigma | AB_10603437 |
| Glutamine synthetase | mouse | Müller glia | 1:1000 | Transduction Laboratories | AB_2313767 |
| GFP | chicken | GFP expressing cells | 1:500 | Aves labs | AB_2307313 |
| PKCβ | mouse | Rod bipolar cells (cytoplasmic) | 1:500 | Santa Cruz Biotechnology | AB_628142 |
| recoverin | rabbit | Photoreceptor (cytoplasmic) + cone bipolar cells (cBPCs) | 1:500 | Millipore | AB_2253622 |
| Red-green opsin | rabbit | LM cones | 1:200 | Millipore | AB_177456 |
| Rhodopsin (Rho4D2) | mouse | Rod photoreceptor | 1:200 | Molday (gift) <sup>38</sup> | N/A |
| Secretagogin | rabbit | Amacrine + cBPC + horizontal cells (cytoplasmic) | 1:200 | Cell Signaling Technology | AB_2798371 |
| β-synuclein | rabbit | Amacrine + ganglion cells (cytoplasmic) | 1:100 | Cell signaling | AB_1904156 |

| Secondary antibodies |  |  |  |  |  |
| --- | --- | --- | --- | --- | --- |
| Conjugate | species | specific for | dilution | supplier | RRID |
| Alexa Fluor 488 | Donkey | Mouse IgG (H+L) | 1:400 | Jackson Immuno Research (West Grove, PA) | AB_2340851 |
| Alexa Fluor 647 | Donkey | Rabbit IgG (H+L) | 1:400 | Jackson Immuno Research | AB_2492288 |

## Results

### Generation of immunodeficient rRosa-LSL-TdTomato rats with retinal degeneration

After injection of 156 pronuclear embryos of RRRC strain #539 with the TdTomato1010 construct, one founder female was identified (out of a litter of 10) (**Figure 1 D-F**). A total of 10 animals were screened for the TdTom-loxP transgene using PCR. Results showed visible band at the expected size of 588bp F1/R1 primer for rat #5, suggesting that it was positive for donor integration (**Figure 1D**). Furthermore, the animal was screened for targeted and random integration, which exhibited positive bands for both at expected size of 1354bp and 750bp, respectively (**Figure 1 E,F**).

The animal was designated as founder, and was backcrossed to rhodopsin-mutant immunodeficient WT (RRRC#539) to generate ten F1 heterozygous progeny at Envigo which were then shipped to UCI and bred to homozygosity. **Figure 3** shows an example of a litter of the F8 generation that was all homozygous.

**Figure 3:**
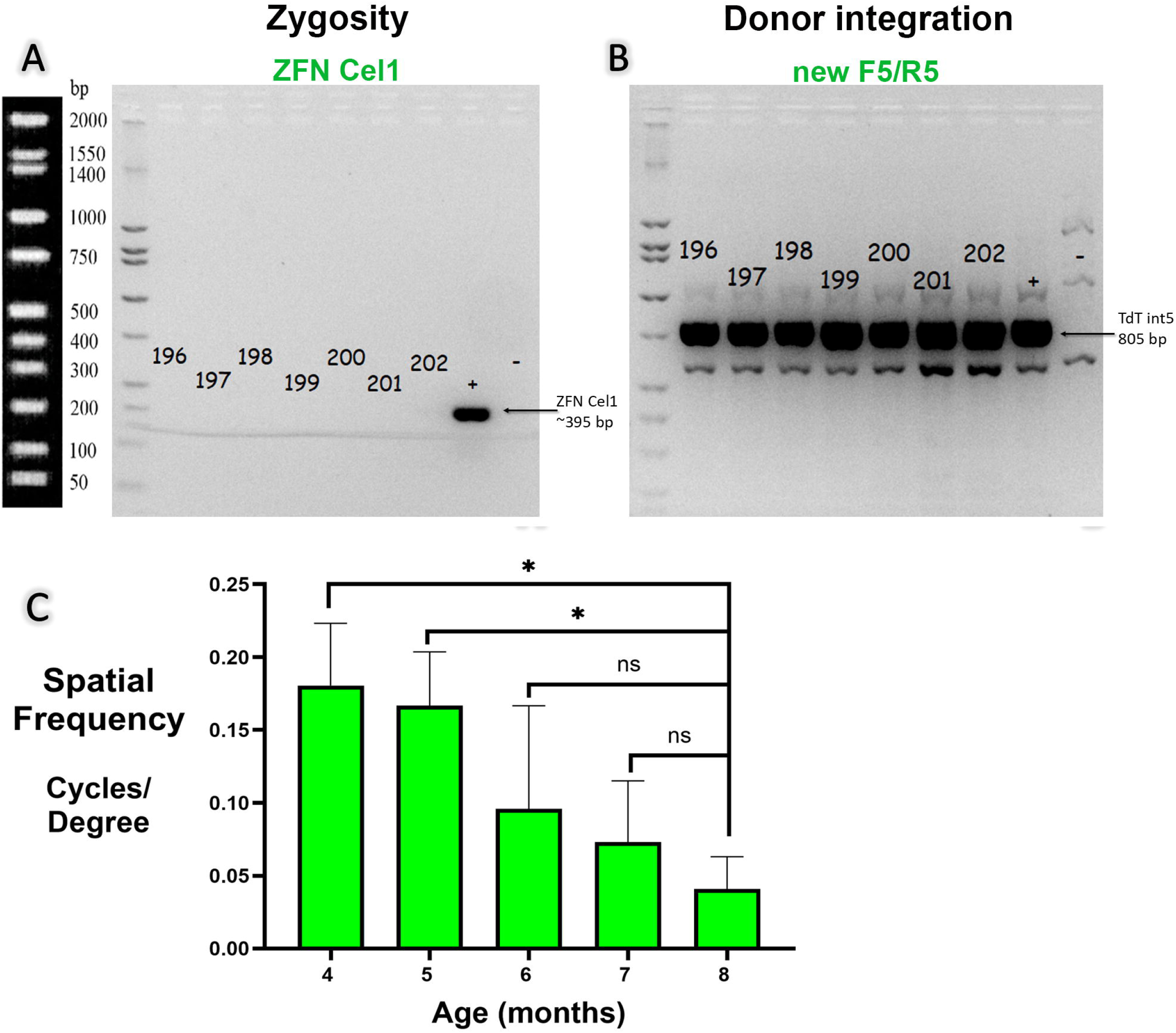
R**a**ts **homozygous for the CAG-LSL-TdTomato transgene.** Genotyping results of TdTomato 1010 litter (F8) generation (all homozygous). **A)** ZFN Cel1 primer for zygosity (only expressed by WT and heterozygotes). **B)** Donor Integration: all pups express transgene (new F5/R5). **C) Optokinetic testing results** between the ages of 4 – 8 months (N=9 for 4 and 5 months of age; N=3 for 6 months of age; N=5 for 7 and 8 months of age). Continuous deterioration of visual acuity. Visual acuity at 8 months is significantly different to visual acuity at 4 and 5 months (unpaired t-test with Welch’s correction, p<0.05).

Gel electrophoresis indicated no visible bands for WT gene screened by ZFN Cel1 primers (**Figure 3A**) and visibly bright bands for the internal transgene screened by TdTom int5 primers (**Figure 3B**) at expected size of 805bp. These results revealed that these rats had successfully obtained the homozygous rRosa-LSL-TdTomato gene sequence. We have submitted this strain (7 nude males, age 71d; and 19 females, age 27-35d) to the RRRC, University of Missouri (future strain RRRC #1055; strain name: *SD-Foxn1rnuTg((Rho-S334X)3,CAG-tdTomato)1010Mjsuc*).

### Visual test of rats with CAG-LSL-TdTomato gene shows vision loss over time

OKT was used to test the vision of rats homozygous for the CAG-LSL-TdTomato gene. As the rats aged, they exhibited a continuous and significant decrease in visual acuity over the age of 4 – 8 months (**Figure 3C**)..

### Generation of Pcp2-Cre rats

Envigo designed sgRNAs targeting the stop codon of rat *Pcp2*, and screened for the most active sgRNAs in cultured cells first, then designed and synthesized donor DNA, containing the specified *Pcp2-Cre* mutation (**Figure 2A-C**). This donor DNA along with Cas9/sgRNA reagent was then delivered into fertilized embryos. However, no transgenic founders could be identified after injecting the *Pcp2-Cre* donor DNA in the immunodeficient *Rho-S334ter-3* rat strain (536 embryos). After switching to Long-Evans rats, 2 founders were identified after injection of 291 embryos, both with targeted gene integration, but one of the founders also with randomized gene integration. A total of 7 animals were screened for the Pcp2-Cre transgene using PCR. Results showed a visible band at the expected size of 697bp for rat #4 and #6, suggesting that they were positive for donor integration (**Figure 2D**). Furthermore, the animal was screened for targeted and random integration, where #4 exhibited positive bands for targeted integration, while #6 exhibited positive bands for both targeted and random at expected size of 1037bp and 750bp, respectively (**Figure 2 E,F**). The F1 generation was received in 2024 and bred to homozygosity in 2025. The current *Pcp2-Cre* rats show both random and targeted gene integration (GI).

The animals were designated as founders and were backcrossed to a Long-Evans and to a NIH nude rat, respectively to generate F1 heterozygous progeny at Envigo, then bred to homozygosity at UCI.

Gel electrophoresis resolved a mix of visible bands and no bands for WT gene screened by Pcp2 Cel1 primers at 502bp, and visibly bright bands for the internal Pcp2 transgene screened by Pcp2 int primers at expected size of ∼650bp. **Figure 4** shows a litter of the F3 generation, all expressing the *Pcp2-Cre* transgene (positive for F1/R1 int primer) (**Figure 4B**). Rats Pcp2-186-195 were homozygous (did not have the Cel1 primer). Rats Pcp2-196-198 were also positive for the Cel1 primer, meaning they were heterozygous (**Figure 4A**). These results revealed that the progeny had produced both heterozygous and homozygous *Pcp2-Cre* gene offspring. Rats with the heterozygous *Pcp2-Cre* gene were sacrificed, while rats with the homozygous gene would be conserved and bred to produce successive offspring with the homozygous *Pcp2-Cre* gene.

**Figure 4:**
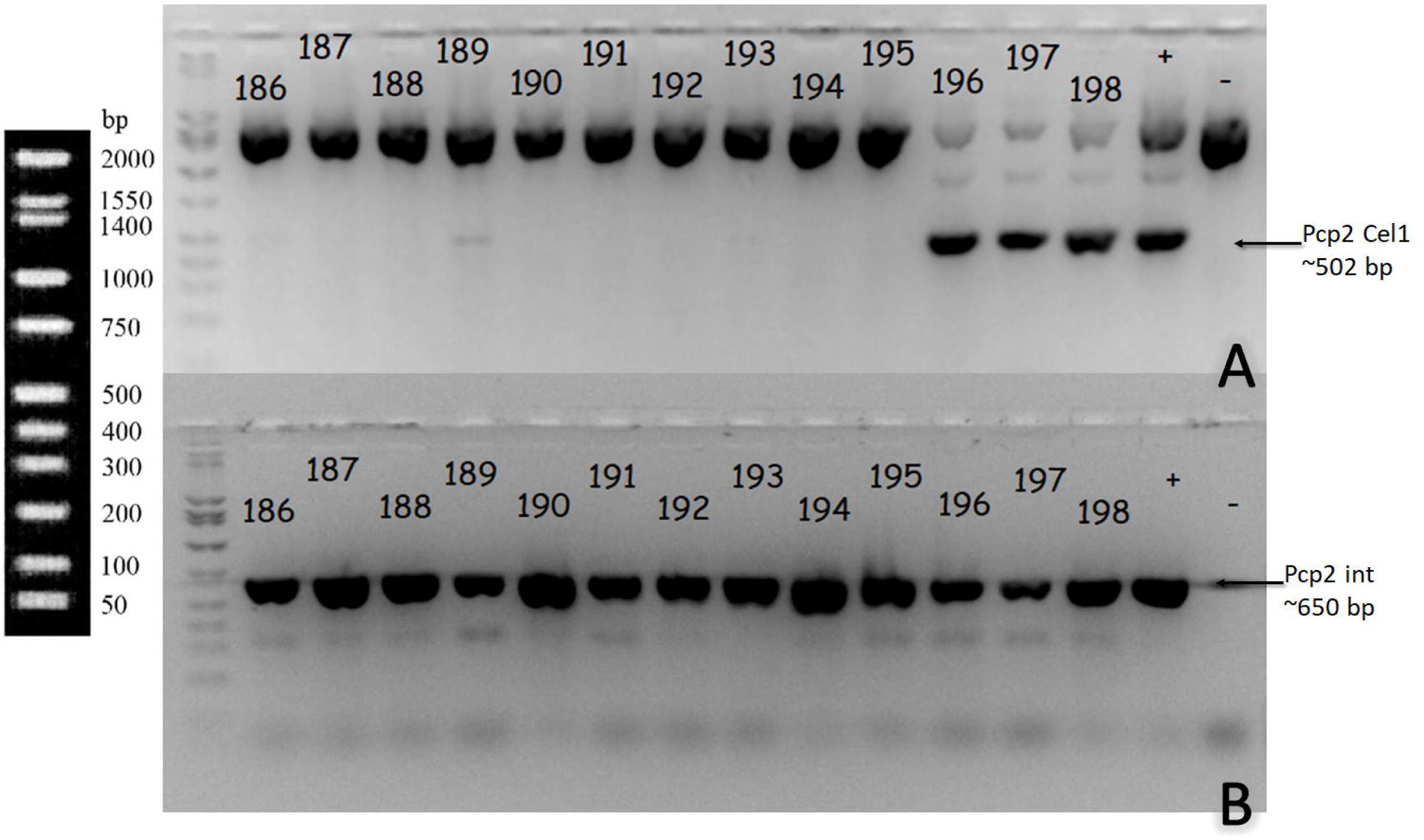
Pcp2-Cre rats of F3 generation **(186-198) t**hat are mixed homo- and **heterozygous for the transgene**. **A)** Pcp2 Cel1 primer shows 3 rats expressing the Pcp2 Cel1 primer (#196-198). **B)** All rats express at least one copy of the Pcp2-Cre transgene as indicated by the Pcp2 F1/R1 primer. This means that rats 196-198 are heterozygous for the transgene.

### F1 offspring of crossing CAG-LSL-TdTomato and Pcp2-Cre rats

Homozygous rats of both strains were mated. All rats of the F1 generation were heterozygous for both transgenes (example in **Figure 5)** and expressed TdTomato (**Figures 6-8**).

**Figure 5:**
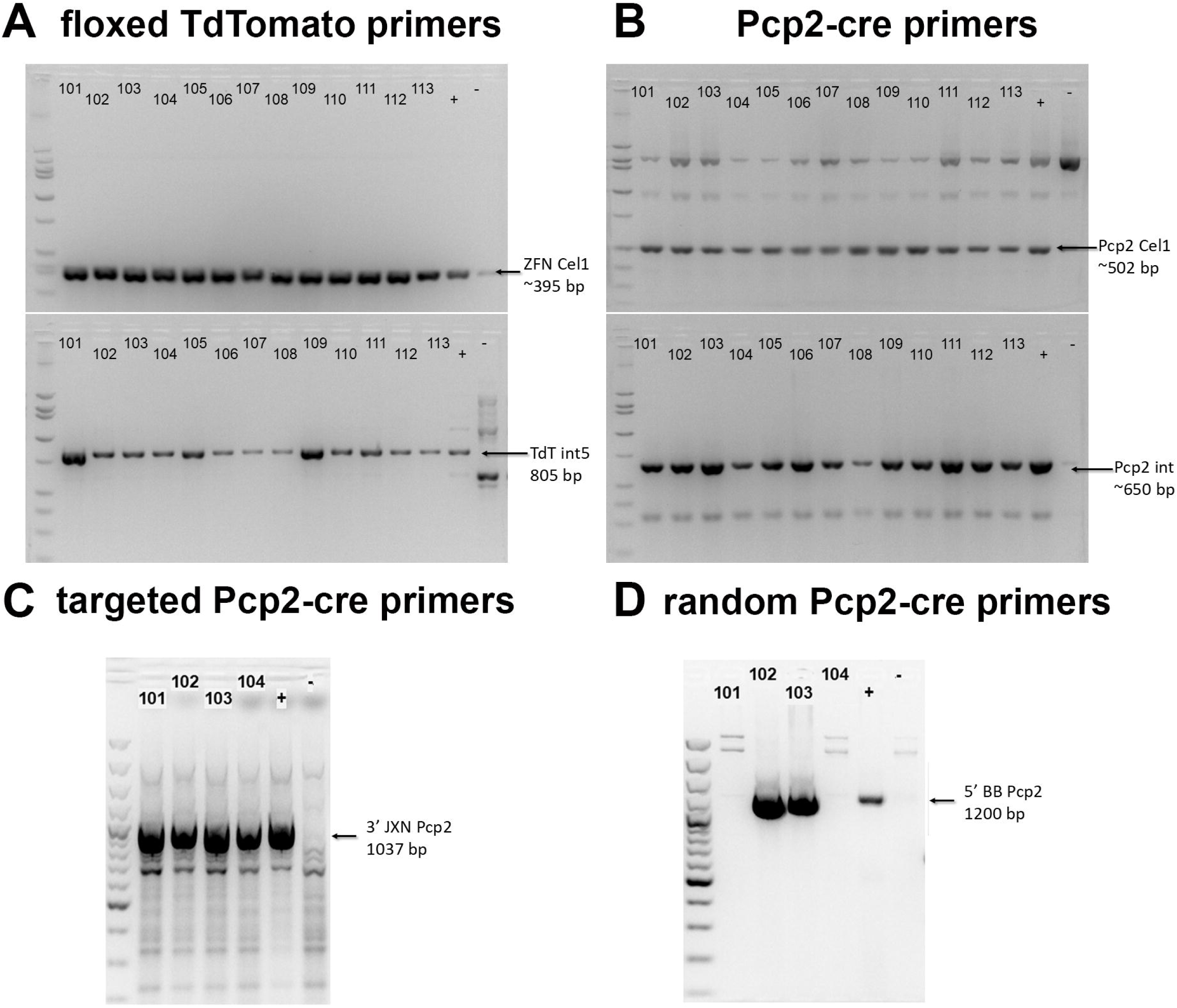
G**e**notyping **results of F1 cross of homozygous CAG-LSL-TdTomato and Pcp2-Cre rats**. All rats are heterozygous for *CAG-LSL-TdTomato* and *Pcp2-Cre*. **A)** Primers ZFN Cel1 (top, for Zygosity, 395 bp) and CAG-LSL-TdTomato transgene (“TdT int5” = new F5/R5, 805 bp). **B)** Primers Pcp2 Cel1(top, for zygosity, 502 bp) and Pcp2 integration (Pcp2 int = F1/R1 primer, 650 bp).

**Figure 6.**
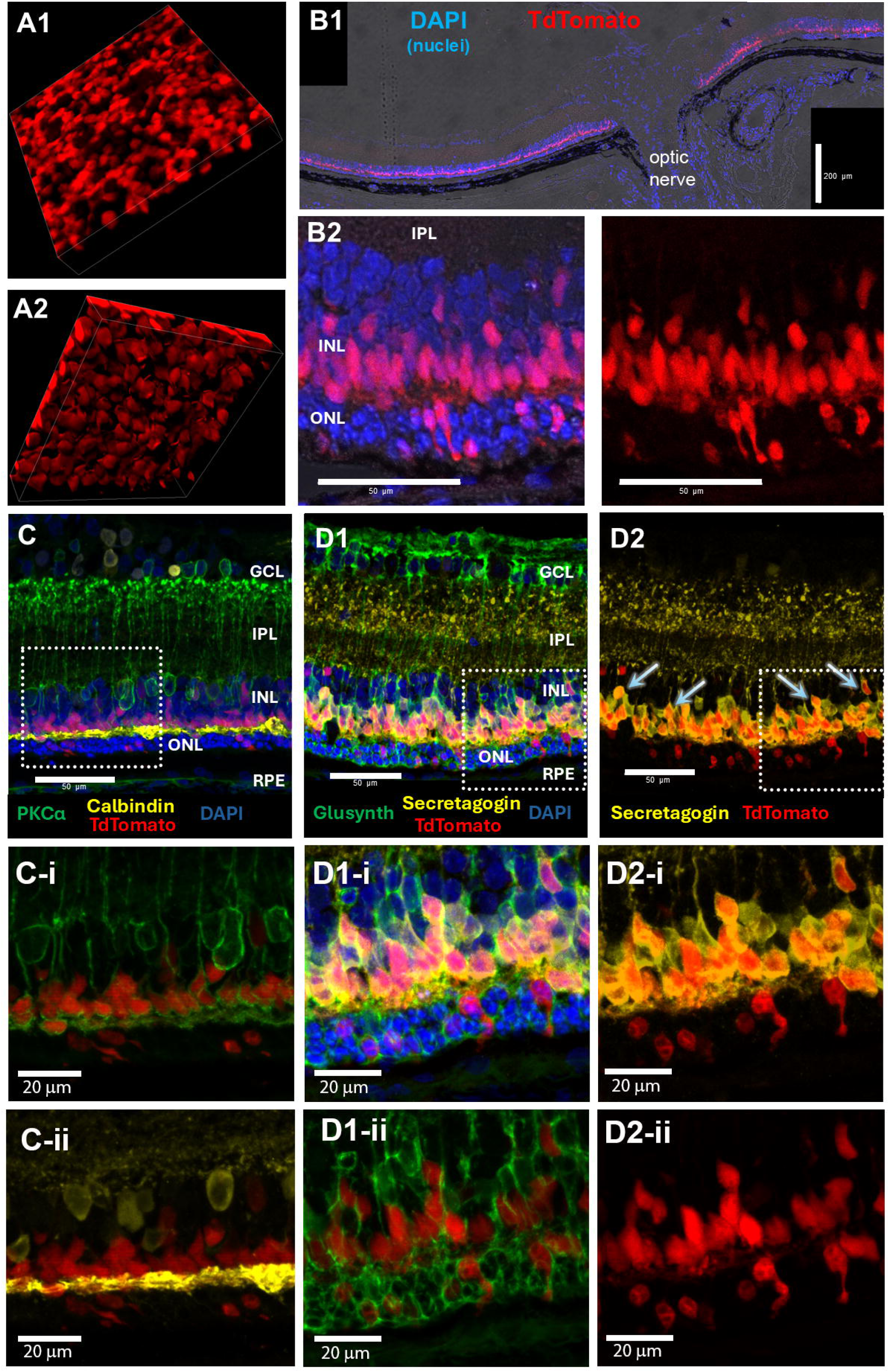
“RTP” rat with targeted TdTomato expression, postnatal day 19. **A)** retinal wholemount stack seen at different angles; **B)** Retinal section through optic disk counterstained with DAPI. **B1)** overview section through optic disk; **B2)** enlargement of different section, showing TdTomato stain in inner nuclear layer and subgroup of photoreceptors. At this age, there is still a thin outer nuclear layer (2-4 nuclei) left. **C)** PKCα (green; rod bipolar cells), Calbindin (yellow; horizontal cells), TdTomato (red). Enlargements: **C-i** (PKCα, TdTomato), **C-ii** (calbindin, TdTomato). Not much overlap between PKCα or Calbindin and TdTomato. **D)** Glutamine synthetase (green; Müller cells), Secretagogin (yellow; cone bipolar cells), TdTomato (red). **D1)** All channels, including DAPI. **D1-i)** enlargement of box in D1. **D1-ii)** Enlargement TdTomato, CRALBP (green). **D2)** Secretagogin (yellow) and TdTomato (red). Overlap between TdTomato and Secretagogin in many cells. **D2-i)** Enlargement of box in D2. **D2-ii**) Enlargement, TdTomato only. Bars = 200µm (B1), = 50µm (B2, C, D1, D2), =20µm (C-i, C-ii, D1-I, D1-ii, D2-I, D2-ii).

### Targeted vs random TdTomato expression

There was a variability of TdTomato expression between the retinal layers and other ocular structures of the eyes. Animals with only targeted integration of the *Pcp2-Cre* transgene (**Figure 6 A1, A2, B1, B2**), expressed TdTomato fluorescent cells specifically in the cone bipolar cells (marker secretagogin) in the inner nuclear layer (INL) (**Figure 6 D1, D2**) in addition to scattered cone photoreceptor cells in the outer nuclear layer (ONL) (**Figure 6**; **Figure 7 E-H**). Interestingly, there was not much co-localization with the rod bipolar cells marker PKCα, but some colocalization with the horizontal cell marker Calbindin (**Figure 6C**). The time course of photoreceptor degeneration between postnatal day (P) 19 – 35 is shown in **Figure 7**, using the markers rhodopsin (**Figure 7A, B**), recoverin (**Figure 7C, D**), red-green opsin (**Figure 7E,F**) and peanut agglutinin (PNA; **Figure 7 G,H**). .The ONL was reduced from 2.5 rows to 1 row between 19 and 35d postnatal. Cones still presented with short outer segments on postnatal day 19 (**Figure 7 E,G**) which looked very distorted and abnormal of postnatal day 35.

**Figure 7.**
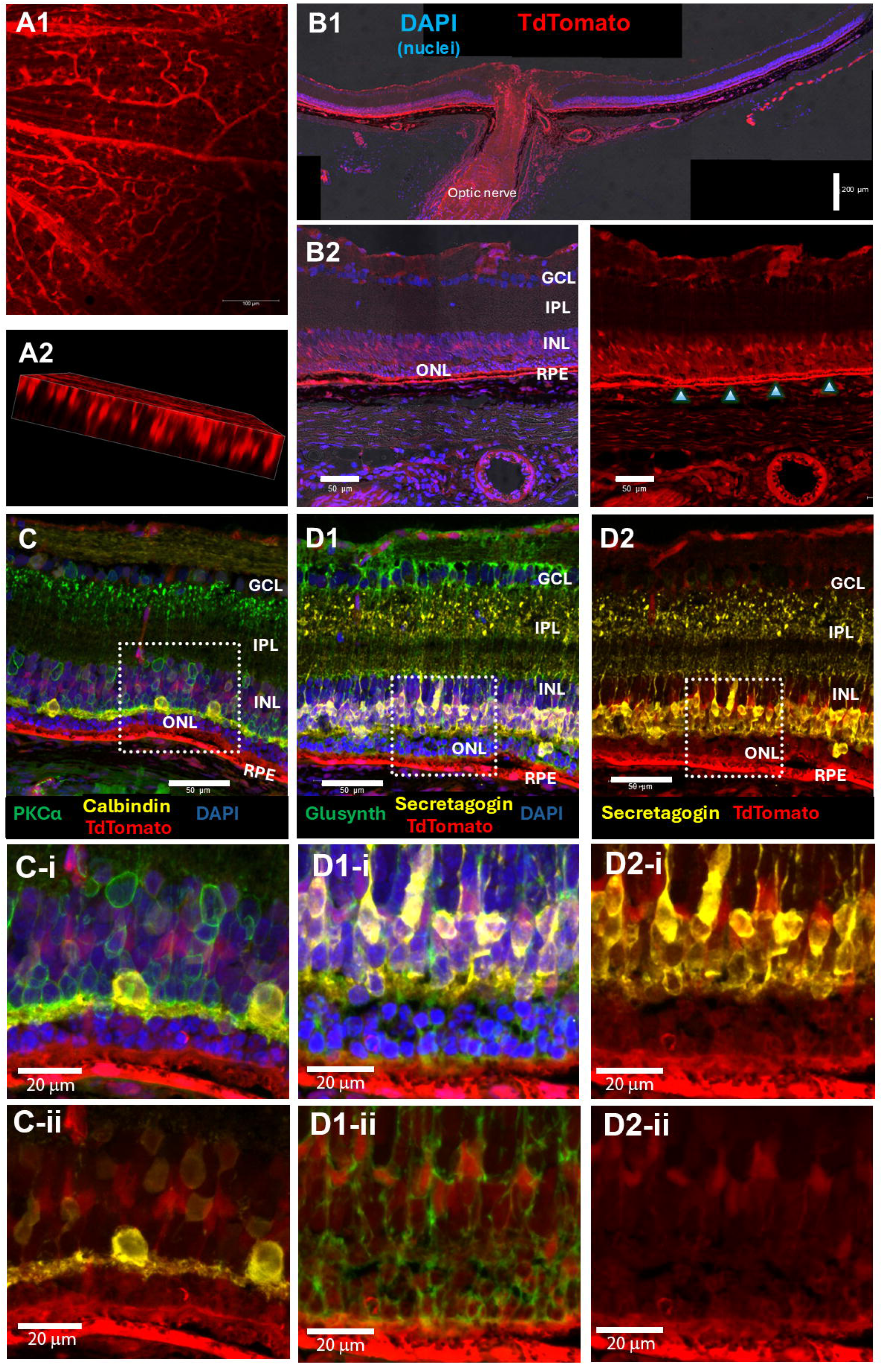
Photoreceptor markers in “RTP” rats with targeted TdTomato expression**show progressive photoreceptor degeneration.** RTP rats with targeted gene insertion, at 2 different ages (postnatal days 19 and 35). All panels show an overview in the top row, and an enlargement (indicated by box) below. **A), B) Rhodopsin** (green). **A)** On d19, about 2 rows of photoreceptors left. Abnormal rhodopsin staining in cell bodies of rods. Few TdTomato cells in ONL. **B)** On d 35, outer nuclear layer is reduced to one row. **C), D) recoverin** (marker for photoreceptor and cone bipolar cells; white) on d19 (C) and d35 (D). **E), F) Cone red-green opsin** staining (white) on d19 (E) and d35 (F). Bars = 50µm (A-H), =20µm (A’-H’). **G, H) Peanut agglutinin** (PNA) staining (marker for cone extracellular matrix, white) on d19 (G) and d35 (H). Note the significant degeneration between 19 and 35 days.

**Figure 8.**
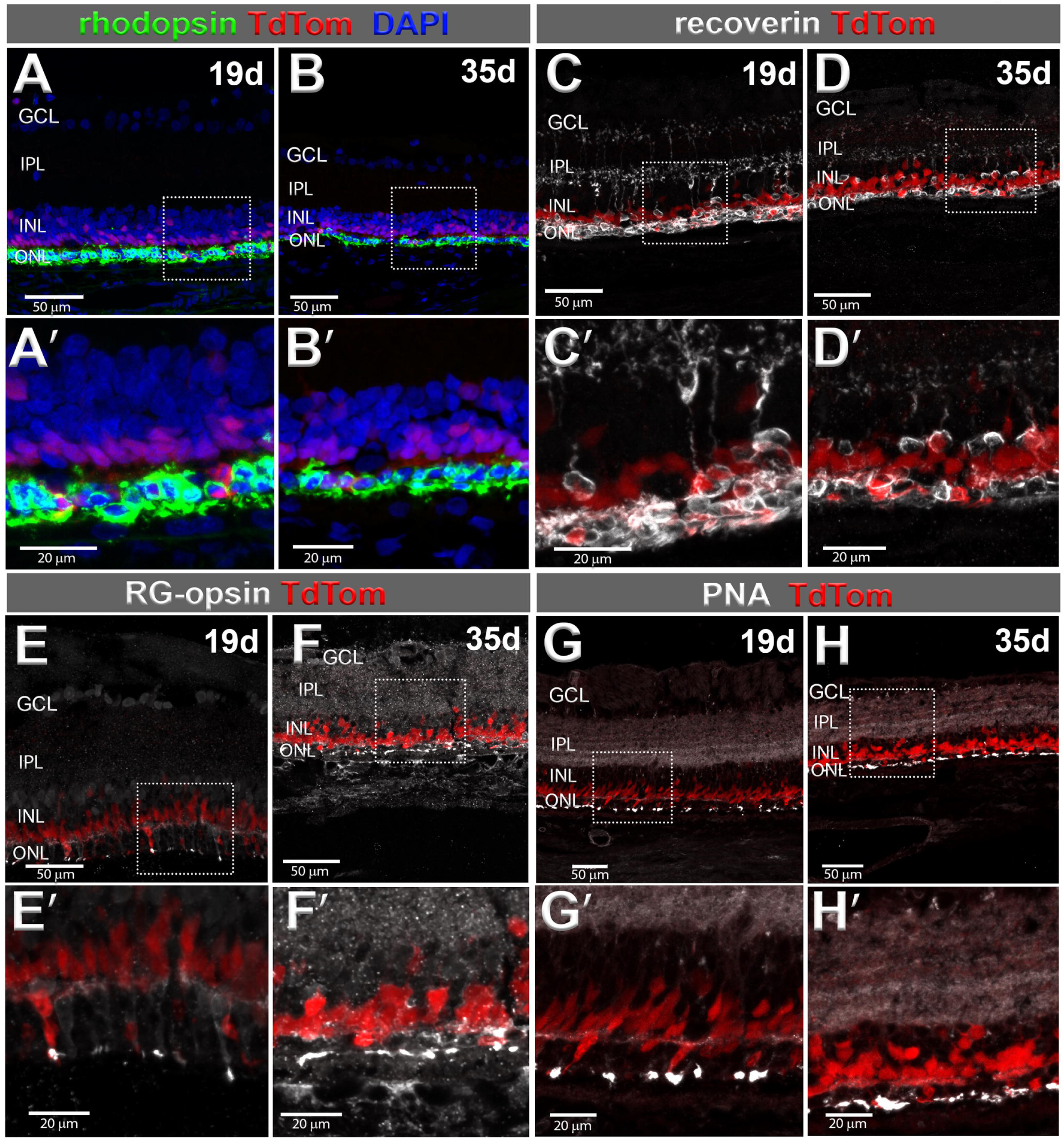
“RTP” rat with random TdTomato expression, postnatal day 19. **A)** retinal wholemount stack seen at different angles. **A1** shows blood vessel staining; **A2** shows radial cells (Müller glia). **B)** Retinal section through optic disk counterstained with DAPI. **B1)** overview section through optic disk (TdTomato stain in optic disk); **B2)** enlargement of different section, showing TdTomato stain in inner nuclear layer, but also in the RPE and in blood vessels. **C)** PKCα (green; rod bipolar cells), Calbindin (white; horizontal cells), TdTomato (red). **C-i** (PKCα, TdTomato), **C-ii** (calbindin, TdTomato). Not much overlap between PKCα or Calbindin and TdTomato. **D)** Glutamine synthetase (green; Müller cells), Secretagogin (white; cone bipolar cells), TdTomato (red). **D1)** All channels, including DAPI. **D1-i)** enlargement of box in D1. **D1-ii)** Enlargement TdTomato, CRALBP (green). **D2)** Secretagogin (white) and TdTomato (red). **D2-i)** Enlargement of box in D2. **D2-ii**) Enlargement, TdTomato only. Overlap between TdTomato and Secretagogin in subgroup of cells. Note additional TdTomato label in RPE, blood vessels and glial cells. Bars = 200µm (B1), = 50µm (B2, C, D1, D2), =20µm (C-i, C-ii, D1-I, D1-ii, D2-I, D2-ii).

As predicted from the genotype of the F1 “RTP” rats, animals containing both targeted and random integrated *Pcp2-Cre* transgene (**Figures 8 - 10**), exhibited both random and targeted TdTomato fluorescent signals in the retinal bipolar cells (BPCs), as well as the RPE, Müller glia and astrocytes and blood vessels (**Figure 8, A,B; Figures 9-10**).

**Figure 9.**
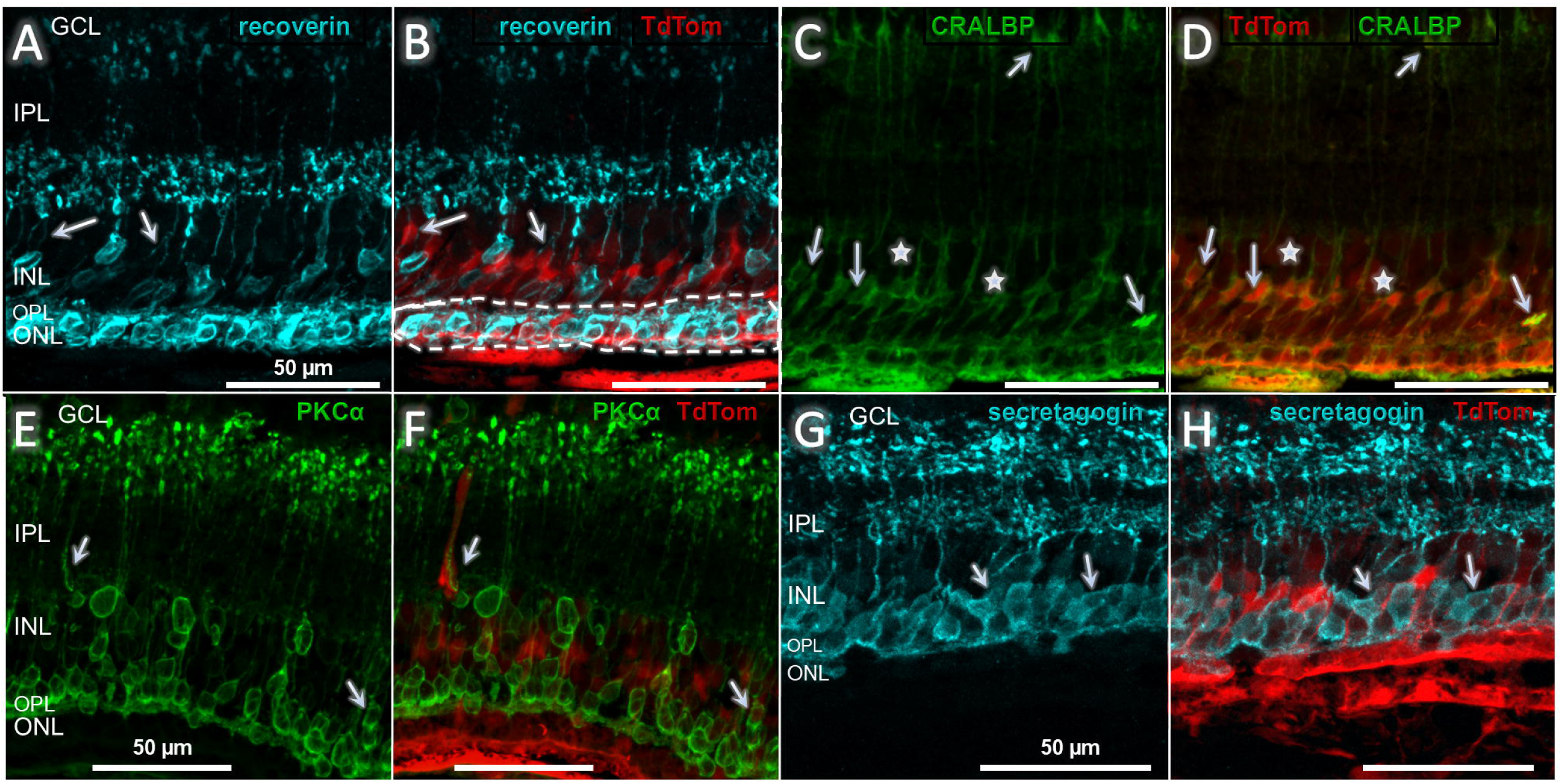
**Staining of “RTP” rats with random TdTomato expression**. Strong TdTomato expression in RPE and radial cells in the retina. **A,B) Recoverin** (turquoise) on d19. Arrow point to same TdTomato+ cells in A) and B). Dashed line in B) indicates photoreceptor layer. **C), D) CRALBP** (green, Müller glia cells) on d19 (same section). Several TdTomato+-cells are co-labeled for CRALBP. **E, F) PKC**α (green, rod bipolar cells). Arrows indicate TdTomato+ cells co-labeled with PKCα. However, TdTomato-structure labeled on left side is a blood vessel in close vicinity to a PKCα-labeled process. **G,H) Secretagogin** (turquoise) on d35. In the random Td-Tomato-expressing rat, there is still relatively faint stain of Secretagogin+ cells, but the stronger label appears in Secretagogin-negative cells (presumably Müller cells. Bars = 50µm.

**Figure 10.**
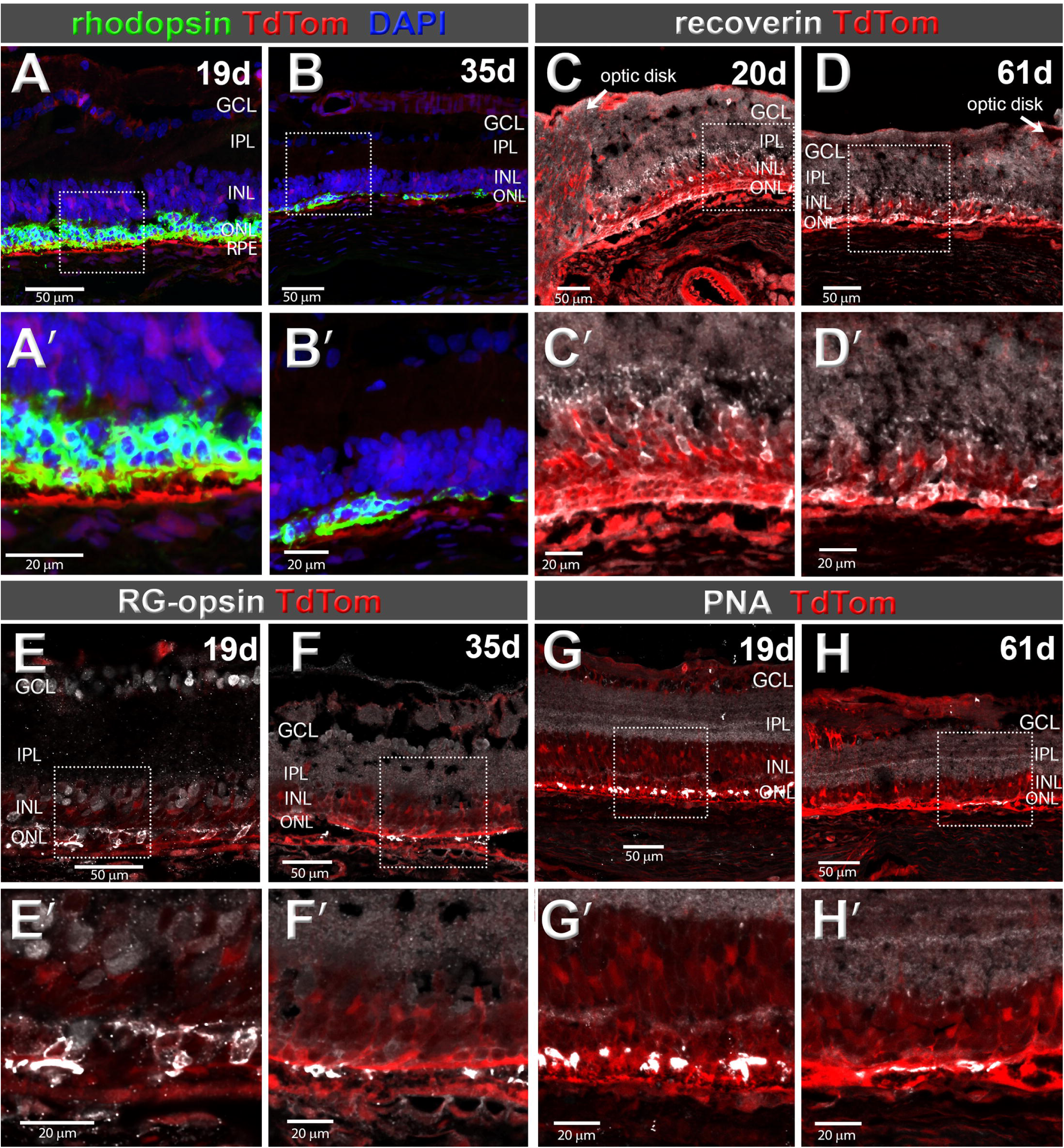
Photoreceptor markers in “RTP” rats with targeted TdTomato expression show progressive photoreceptor degeneration. RTP rats with targeted gene insertion, at 2 different ages (postnatal days 19 and 35). All panels show an overview in the top row, and an enlargement (indicated by box) below. **A), B) Rhodopsin** (green). **A)** On d19, about 2 rows of photoreceptors left. Abnormal rhodopsin staining in cell bodies of rods. Few TdTomato cells in ONL. **B)** On d 35, outer nuclear layer is reduced to one row. **C), D) recoverin** (marker for photoreceptor and cone bipolar cells; white) on d19 **(C)** and d35 **(D)**. **E), F) Cone red-green opsin** staining (white) on d19 **(E)** and d35 **(F)**. Bars = 50µm (A-H), =20µm (A’-H’). **G, H) Peanut agglutinin** (PNA) staining (marker for cone extracellular matrix, white) on d19 **(G)** and d35 **(H)**. Note the extensive photoreceptor degeneration at the age of 61 days.

In animals that expressed both random and targeted integration of the transgene, a variety of retinal cells from different retinal layers expressed the TdTomato signal along with retinal BPCs. Colocalization of the cytoplasmically stained cells (recoverin) in the ONL with TdTomato-suggested that some photoreceptors also arbitrarily expressed the fluorescent signal, while, at the same time, recoverin-positive cone BPCs from the INL also showed colocalization with the TdTomato-positive cells (**Figure 9A,B**; **Figure 10 C,D**). Further examination of the INL also showed that the TdTomato signal colocalized with Glutamine synthetase (**Figure 8D**) and CRALBP (Muller glia) (**Figure 9C,D**), PKC[zl (rBPCs) (**Figure 9 E,F**), and secretagogin (cBPCs) (**Figure 8D**), which suggested that these retinal cells were also affected by the expression of TdTomato. Photoreceptor degeneration between P19 and P61 was similar in rats with both targeted and random *Pcp2-cre* expression (**Figure 10**).

### Examples of pilot GFP transplants in TdTomato host

Figure 11 shows examples of GFP-expressing rat retinal transplants to “RTP” rats with targeted or targeted/random *Pcp2-Cre* expression. The transplant to a rat with targeted *Pcp2-Cre* expression in Figure 11 **A,B** (37 days post-surgery = dps), contained photoreceptors in a rosette, showed intermingling to transplant and host cells at the transplant host interface, extension of transplant processes into the host INL, and extension of host processes into the transplant. A transplant to a rat with random *Pcp2-Cre* expression (Figure 11 **C,D**, 77dps) had developed a photoreceptor layer with outer segments in contact with host RPE expressing TdTomato. The enlargement showed host cell processes, as well as host-derived blood vessels extending into the transplant. Donor cells had migrated into the host retina (**Figure 11C**).

**Figure 11.**
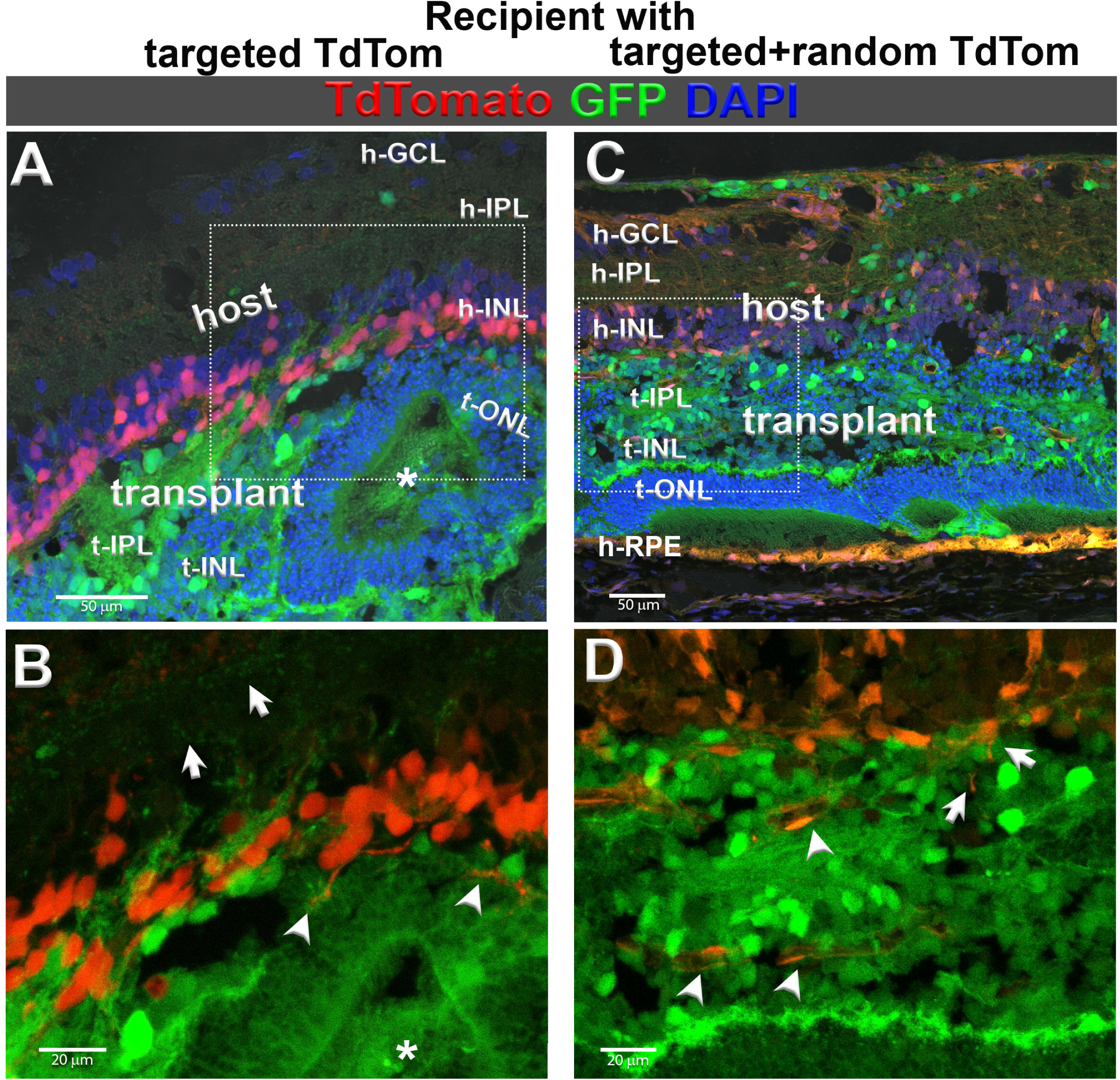
Td-Tomato-labeled *RhoS334ter-3* recipients with transplants of GFP-expressing rat retinal sheets (grafted on embryonic day E19). Sections stained with antibody against EGFP. Top (A, C): TdTomato, GFP, and DAPI (nuclei), boxes indicate areas of enlargement in bottom (B, D) shown with red and green channels only. **A) B):** recipient with targeted TdTomato expression (as described in Figs. 6, 8). **A)** Transplant with photoreceptor rosette, lumen indicated by asterisk. Arrows in the enlargement in **B)** point out GFP+ processes in the host inner plexiform layer. Arrowheads point to host processes extending into the transplant. **C), D)**: recipient with random TdTomato expression in a variety of cell types (as described in Figures 7, 9, 10). **C)** Transplant has developed outer nuclear layer, with photoreceptor outer segments towards the host RPE expressing TdTomato. **D)** Enlargement. Arrowheads point to host-derived, TdTomato expressing blood vessels in transplant. Arrows point to cell processes extending into the transplant. Bar in A, C) = 50µm; Bars in B, D) = 20µm

## Discussion

The *Cre-loxP* system is universally useful to create cell type-specific gene expression or label (review ^30, 31^). In the current study, we have generated immunodeficient retinal degenerate *RhoS334ter-3* rats expressing a CAG-LSL-TdTomato transgene (*TdTomato1010*, “RNT”, future RRRC strain #1055), When crossed to cre-expressing strains, the F1 offspring will express TdTomato in *Cre*-expressing cells (“RTP”) and show severe retinal degeneration as the rhodopsin mutation is dominant. We have created immunodeficient *Pcp2-Cre* -expressing rats from two founders, both with targeted gene insertion. However, the second founder expressed *Pcp2-Cre* also randomly.

In contrast to *Pcp2-Cre* mouse models with specific label of rod bipolar cells ^24, 25^, our rat model with targeted Pcp2-Cre expression showed most of the TdTomato label in cone bipolar cells, not in rod bipolar cells, and in some additional cone photoreceptors. Thus, the function of the *Pcp2* gene may be different between mice and rats.

The creation of this new rat strain can be beneficial to studying retinal degeneration in several ways. The parents, containing the *Foxn1* gene mutation, as well as the *Pcp2-Cre* and *TdTomato-LoxP*, have produced a generation of a novel hybrid model, where the descendants contain both the *Foxn1* mutation and TdTomato-fluorescent *Pcp2* gene in the retina. Rats with the *Foxn1* gene mutation can be bred to express either wild type (*Foxn1^+/-^*), which produces immunocompetence that can be useful for studying allo-transplantation in the rat model, or homozygous recessive (*Foxn^-/-^*), which exhibits phenotypically immunodeficient characteristics that is suitable for xenotransplantation of the human embryonic stem cell-derived retinal organoid (hESC-RO) to minimize immune rejection. On the other hand, the creation of a cell-specific fluorescent labelling, such as of retinal bipolar cells, has proven the effectiveness and versatility of the *Cre-LoxP* system. Moreover, retinal bipolar cells are an important structure of the retina, which play an important role in initial signal integration and processing before passing onto high-order visual processing in the central nervous system, and their ability to adapt in retinal degeneration is an attractive field of vision research. The novel hybrid model, therefore, offers a more comprehensive versatility to further elucidate the molecular basis of retinal degeneration and how it affects the later components of the retinal circuit, as well as understand the feasibility of hESC-RO in rescuing or delaying the prognosis of retinal degeneration.

Using the ZFN Cel-1 method, rats were able to acquire the TdTomato gene flanked between the loxP sites. Despite its high affinity, there was still a mix of random and integrated transgene in the animals. However, the effect of random integration of the *loxP* gene was not as severe as seen in that of the *Cre* gene. *CRISPR-Cas9* technology, in spite of its versatility and efficiency, was employed to generate a mixed batch of random and targeted integration. Because the *Cre* protein was responsible for dictating which cells would express the TdTomato fluorescent gene, random integration of the *Cre* gene sequence outside of the *Pcp2* promoter site would lead to random TdTomato expression of other cells besides *Pcp2*-expressing cells, as seen in the immunofluorescent results. According to our results, certain ocular structures, such as the RPE, photoreceptors, Müller cells, retinal ganglion cells, and retinal blood vessels, exhibited much stronger TdTomato expression than retinal bipolar cells in the rats expressing random *Pcp2-Cre*. Some rats also appeared to have pink skin, hair, nails, and lens; therefore, depending on the location of the *Cre* transgene, results may vary, and would not be useful to study specific retinal cell structure, such as the bipolar cells. Therefore, rats should be examined for phenotypically abnormal appearance (pink hair, skin, lens, and transparent RPE) coupled with genotyping to eliminate random TdTomato expression.

There are several issues that make it more difficult to create transgenic rats versus mice While mouse embryonic stem (ES) cells have been well-characterized and used since the 1980s for gene targeting, stable and pluripotent rat ES cell lines have historically been more difficult to derive and maintain ^32^. This has limited the use of homologous recombination for targeted genetic manipulation in rats. Rats have more complex reproductive physiology, making techniques like pronuclear injection and embryo manipulation technically harder and less efficient compared to mice. Rat embryos are more fragile and sensitive to manipulation, which leads to lower survival and integration rates of the foreign DNA. The development and application of gene-editing tools like CRISPR/Cas9 and ZFNs in rats occurred later than in mice. Though this has improved transgenic rat generation in recent years, mice still remain the more accessible and widely used model.

Despite these challenges, transgenic rats have remained a favorable animal model for *in vivo* study of human diseases such as retinitis pigmentosa and macular degeneration. Rats possess larger eyes than mice, making them more suitable for ophthalmic intervention and retinal imaging. Rats can also model specific mutations associated with human retinal diseases, making it a valuable tool for *in vivo* testing of gene therapy, stem cell replacement, and molecular pathways of the visual circuit. Moreover, rats have a better attention span compared to mice, which guarantee them superior candidates for complex visual and cognitive tasks necessary for behavioral assessment. Due to the advantage of size, rats allow more complex surgical procedures and sampling of blood and tissues, providing a more comprehensive testing of certain diseases. With these reasons, transgenic rat models have proven their unparalleled contribution to vision research in humans.

In addition to our developed *Pcp2-TdTomato* rat model, the CAG-LSL-TdTomato immunodeficient RD rats (future RRRC strain #1055) can be crossed with other *Cre*-expressing rat strains available from the RRRC (Rat Research Resource Center, University of Missouri), e.g. *ChAT-Cre* ^33^, *Pvalb-Cre* ^34, 35^ or *Gad1*-Cre . Alternatively, rats can be injected intravitreally with viruses containing Cre in combination with cell-type specific promoters (e.g. *CamKII, c-fos* or *synapsin* ^31, 36, 37^) which will be useful to study connectivity of human stem-cell derived retinal or photoreceptor transplants. We have submitted this strain to the RRRC so it will be available for other researchers studying retinal transplantation and retinal degeneration. Strain cryopreservation has now been completed.

## Supporting information

1. Donor Design_rRosa-tdTomato-1010

2. Donor Design_ Pcp2-1105 donor DNA.pdf

## Acknowledgments

Supported by NIH R01EY031834, R01EY032948. This work was made possible, in part, through access to the Optical Biology Core Facility of the Developmental Biology Center, a shared resource supported by the Cancer Center Support Grant (CA-62203) and Center for Complex Biological Systems Support Grant (GM-076516) at the University of California, Irvine. We acknowledge support from NIH core grant P30 EY034070 and from an unrestricted grant from Research to Prevent Blindness to the Gavin Herbert Eye Institute at the University of California, Irvine.

We wish to thank the UCI Transgenic Mouse facility for help with genotyping of the *foxn1* mutation. We appreciate the technical assistance from staff and students of the Seiler lab: Huong Do, Akashi Suon, Robert Sims, Meredith Liu, John Boutros, Quinn Tran, Jack Cho, Fiaza Ali, Irene Jin, Thuy-Lin Tran, Yuhan Chen, Umer Shahab, Jack Wang, Nevin V. Joseph, Cassidy Azores, and Renee Mercado.

## Abbreviations

bp: base pairs
BPC: Bipolar cell
CAG: Strong, synthetic and ubiquitous promoter
CFBS: carbo-free blocking solution (for lectin staining)
CRALBP: cellular retinaldehyde binding protein (marker for RPE and Müller glia)
Cre: Cre-recombinase
CRISPR-Cas9: “clustered regularly interspaced short palindromic repeats” genetic engineering, based on bacterial antiviral defense system
DAPI: 4’6’-diamidino-2-phenylinode hydrochloride (nuclear stain)
dps: days post-surgery
DSB: DNA double strand breaks
GCL: Ganglion cell layer
GFP: Green fluorescent protein
gRNA: Guide RNA
h-: host-
hESC: Human embryonic stem cells
HR: High fidelity homologous recombination
INL: Inner nuclear layer
IPL: Inner plexiform layer
LE: Long-Evans rat
loxP: locus of X-over P1, specific DNA sequence recognized by cre recombinase
NHEJ: Non-homologous end joining
ONL: Outer nuclear layer
P: Postnatal day
Pcp2: Purkinje cell protein 2 (L7)
PKCα PNA: Protein kinase C alpha (marker for rod bipolar cells) Peanut agglutinin (marker for cone interphotoreceptor matrix)
RD: Retinal degeneration, retinal degenerate
“RNT”: Retinal degenerate rats expressing CAG-LSL-TdTomato
RO: Retinal organoid
RP: Retinitis pigmentosa
RPE: Retinal pigment epithelium
RPM: Rotations per minute (centrifuge)
rROSA: Genomic safe harbor location
RRRC: Rat Research and Resource Center , MO
“RTP”: Cross of TdTomato1010 with Pcp2-Cre rats
sgRNA: Single guide RNA
t: transplant-
TdTom: TdTomato
UCI: University of California, Irvine
WT: Wildtype
ZFN: Zinc finger nuclease

## Commercial Relationships Disclosure

Magdalene Seiler: Commercial Relationship: Code N (No Commercial Relationship); Helios Nguyen: Commercial Relationship: Code N (No Commercial Relationship); Bin Lin: Commercial Relationship: Code N (No Commercial Relationship); Devan Endejan: Commercial Relationship(s); Inotiv: Code E (Employment): Inotiv; Guojun Zhao: Commercial Relationship(s); Inotiv: Code E (Employment): Inotiv; Lauren Klaskala: Commercial Relationship(s); Inotiv: Code E (Employment): Inotiv

## Author Contribution Statement

Designed the experiment: MJS, BL, GZ, LK Conducted the experiment: HN, DE, BL, GZ Analyzed/interpreted data: MJS, HN, DE, BL, GZ, LK Provided materials: MJS, LK

Wrote the article: MJS, HN Proofed/revised article: MJS, HN, BL, GZ, LK

## Supplemental files

1. Donor Design_rRosa-tdTomato-1010.docx

2. Donor Design_ Pcp2-1105 donor DNA.docx

## References

1. Suleman N. Current understanding on Retinitis Pigmentosa: a literature review. Front Ophthalmol (Lausanne) 2025;5:1600283.

2. Kamde SP, Anjankar A. Retinitis Pigmentosa: Pathogenesis, Diagnostic Findings, and Treatment. Cureus 2023;15:e48006.

3. Fleckenstein M, Schmitz-Valckenberg S, Chakravarthy U. Age-Related Macular Degeneration: A Review. JAMA 2024;331:147–157.

4. Guymer RH, Campbell TG. Age-related macular degeneration. Lancet 2023;401:1459–1472.

5. Confalonieri F, La Rosa A, Ottonelli G, et al. Retinitis Pigmentosa and Therapeutic Approaches: A Systematic Review. J Clin Med 2024;13.

6. Yang J, Lewis GP, Hsiang CH, et al. Amelioration of Photoreceptor Degeneration by Intravitreal Transplantation of Retinal Progenitor Cells in Rats. Int J Mol Sci 2024;25.

7. Thomas BB, Rajendran Nair DS, Rahimian M, Hassan AK, Tran TL, Seiler MJ. Animal models for the evaluation of retinal stem cell therapies. Prog Retin Eye Res 2025;106:101356.

8. LaVail MM, Nishikawa S, Steinberg RH, et al. Phenotypic characterization of P23H and S334ter rhodopsin transgenic rat models of inherited retinal degeneration. Exp Eye Res 2018;167:56–90.

9. Thomas BB, Zhu D, Lin TC, et al. A new immunodeficient retinal dystrophic rat model for transplantation studies using human-derived cells. Graefes Arch Clin Exp Ophthalmol 2018;256:2113–2125.

10. Seiler MJ, Aramant RB, Jones MK, Ferguson DL, Bryda EC, Keirstead HS. A new immunodeficient pigmented retinal degenerate rat strain to study transplantation of human cells without immunosuppression. Graefes Arch Clin Exp Ophthalmol 2014;252:1079–1092.

11. Ishikura M, Muraoka Y, Hirami Y, Tu HY, Mandai M. Adaptive Optics Optical Coherence Tomography Analysis of Induced Pluripotent Stem Cell-Derived Retinal Organoid Transplantation in Retinitis Pigmentosa. Cureus 2024;16:e64962.

12. Liu YV, Santiago CP, Sogunro A, et al. Single-cell transcriptome analysis of xenotransplanted human retinal organoids defines two migratory cell populations of nonretinal origin. Stem Cell Reports 2023;18:1138–1154.

13. Watari K, Yamasaki S, Tu HY, et al. Self-organization, quality control, and preclinical studies of human iPSC-derived retinal sheets for tissue-transplantation therapy. Commun Biol 2023;6:164.

14. Tu HY, Watanabe T, Shirai H, et al. Medium- to long-term survival and functional examination of human iPSC-derived retinas in rat and primate models of retinal degeneration. EBioMedicine 2019;39:562–574.

15. Sims R, Lin B, Xue Y, et al. Effect of immunosuppression on hESC-derived retina organoids in vitro and in vivo. Stem Cell Res Ther 2025;16:165.

16. Lin B, Singh RK, Seiler MJ, Nasonkin IO. Survival and Functional Integration of Human Embryonic Stem Cell-Derived Retinal Organoids After Shipping and Transplantation into Retinal Degeneration Rats. Stem Cells Dev 2024;33:201–213.

17. Thomas BB, Lin B, Martinez-Camarillo J-C, et al. Co-grafts of Human Embryonic Stem Cell Derived Retina Organoids and Retinal Pigment Epithelium for Retinal Reconstruction in Immunodeficient Retinal Degenerate Royal College of Surgeons Rats. Frontiers in Neuroscience 2021;15:752958.

18. Lin B, McLelland BT, Aramant RB, et al. Retina Organoid Transplants Develop Photoreceptors and Improve Visual Function in RCS Rats With RPE Dysfunction. IOVS 2020;61:34.

19. McLelland BT, Lin B, Mathur A, et al. Transplanted hESC-Derived Retina Organoid Sheets Differentiate, Integrate, and Improve Visual Function in Retinal Degenerate Rats. IOVS 2018;59:2586–2603.

20. Xue Y, Lin B, Chen JT, Tang WC, Browne AW, Seiler MJ. The Prospects for Retinal Organoids in Treatment of Retinal Diseases. Asia Pac J Ophthalmol (Phila) 2022;11:314–327.

21. Malhotra S, Seiler MJ, Browne AW. Challenges and Advances in the Production of Transplantable Retinal Tissue from Retinal Organoids. J Ophthalmic Vis Res 2025;20.

22. Cowan CS, Renner M, De Gennaro M, et al. Cell Types of the Human Retina and Its Organoids at Single-Cell Resolution. Cell 2020;182:1623–1640 e1634.

23. Yamasaki S, Tu HY, Matsuyama T, et al. A Genetic modification that reduces ON-bipolar cells in hESC-derived retinas enhances functional integration after transplantation. iScience 2022;25:103657.

24. Zhang XM, Chen BY, Ng AH, et al. Transgenic mice expressing Cre-recombinase specifically in retinal rod bipolar neurons. IOVS 2005;46:3515–3520.

25. Lu Q, Ivanova E, Ganjawala TH, Pan ZH. Cre-mediated recombination efficiency and transgene expression patterns of three retinal bipolar cell-expressing Cre transgenic mouse lines. Mol Vis 2013;19:1310–1320.

26. Lois C, Hong EJ, Pease S, Brown EJ, Baltimore D. Germline transmission and tissue-specific expression of transgenes delivered by lentiviral vectors. Science 2002;295:868–872.

27. Seiler MJ, Lin RE, McLelland BT, et al. Vision Recovery and Connectivity by Fetal Retinal Sheet Transplantation in an Immunodeficient Retinal Degenerate Rat Model. IOVS 2017;58:614–630.

28. Prusky GT, West PW, Douglas RM. Behavioral assessment of visual acuity in mice and rats. Vision research 2000;40:2201–2209.

29. Douglas RM, Alam NM, Silver BD, McGill TJ, Tschetter WW, Prusky GT. Independent visual threshold measurements in the two eyes of freely moving rats and mice using a virtual-reality optokinetic system. Vis Neurosci 2005;22:677-684.

30. Kim H, Kim M, Im SK, Fang S. Mouse Cre-LoxP system: general principles to determine tissue-specific roles of target genes. Lab Anim Res 2018;34:147–159.

31. Shcholok T, Eftekharpour E. Cre-recombinase systems for induction of neuron-specific knockout models: a guide for biomedical researchers. Neural Regen Res 2023;18:273–279.

32. Kawaharada K, Kawamata M, Ochiya T. Rat embryonic stem cells create new era in development of genetically manipulated rat models. World J Stem Cells 2015;7:1054–1063.

33. Witten IB, Steinberg EE, Lee SY, et al. Recombinase-driver rat lines: tools, techniques, and optogenetic application to dopamine-mediated reinforcement. Neuron 2011;72:721–733.

34. Wright AM, Zapata A, Hoffman AF, et al. Effects of Withdrawal from Cocaine Self-Administration on Rat Orbitofrontal Cortex Parvalbumin Neurons Expressing Cre recombinase: Sex-Dependent Changes in Neuronal Function and Unaltered Serotonin Signaling. eNeuro 2021;8.

35. Patrono E, Hruzova K, Svoboda J, Stuchlik A. The role of optogenetic stimulations of parvalbumin-positive interneurons in the prefrontal cortex and the ventral hippocampus on an acute MK-801 model of schizophrenia-like cognitive inflexibility. Schizophr Res 2023;252:198–205.

36. Kawabata H, Konno A, Matsuzaki Y, et al. Improving cell-specific recombination using AAV vectors in the murine CNS by capsid and expression cassette optimization. Mol Ther Methods Clin Dev 2024;32:101185.

37. Juttner J, Szabo A, Gross-Scherf B, et al. Targeting neuronal and glial cell types with synthetic promoter AAVs in mice, non-human primates and humans. Nat Neurosci 2019;22:1345–1356.

38. Molday RS, MacKenzie D. Monoclonal antibodies to rhodopsin: characterization, cross-reactivity, and application as structural probes. Biochemistry 1983;22:653–660.

