## Supplementary material for "Generation of a new immunodeficient rat model of retinal degeneration with LSL TdTomato reporter and TdTomato-Pcp2 expression": 1. Donor Design_rRosa-tdTomato-1010

**Donor DNA Design**

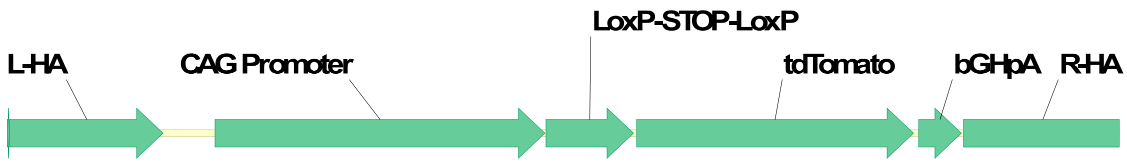


**Sequence (5733bp):**

Ccgcgtgtgaaaacacaaatggcgtgttttggttggagtgaggcgcctgtcaattaacggctgccggagtgcgcagccgctgactgcctcgctgtgcc
cactgggtggggcgggaggtaggtggggtgaggcgagctggacgtgcgggcgcggtcggcctctggcggggcgggggaggggagggtcagcgaaa
gtggctggcgcgtgagcggcctcccaccctcccсttcctctgggggagtcgttttacccgccgccggcctggcctcgtcatctgattggctctcggggctc
agaaaactggcctttgcaattggcccgcgttcatgcaagttcagtccctaagctggctggcgggggcggcagggaggcgctcacaggttccggccctc
cccccaggccccgcgccgcagagtctggccccgcgcccctgcgcaacgtggcaggaagcgcgcgctgggggcggggacgggcggtcggtctgagcg
gcgggcgggtgcaaacgggattcctccttgagttgtggcactgaggaacgtgctgaacaagacctacattgcactccagggagtggatgaaggagtt
ggggctcagtcgggttgtattggagacaagaagcacttgctctccaaaagtcggtttgagttatcattaagggagctgcagtggagtaggcggagaaa
aggccgcacccttctcaggacgggggaggggagtgttgcaatacctttctgggagttctctgctgcctcctgtcttctgaggaccgccctgggcctggaa
gattcccttccccggatccttggggttgcgccttttccaaggcagccctgggtttgcgcagggacgcggctgctctgggcgtggttccgggaaacgcag
cggcgccgaccctgggtctcgcacattcttcacgtccgttcgcagcgtcacccggatcttcgccgctacccttgtgggccccccggcgacgcttcctgctc
cgcccctaagtcgggaaggttccttgcggttcgcggcgtgccggacgtgacaaacggaagccgcacgtctcactagttattaatagtaatcaattacg
gggtcattagttcatagcccatatatggagttccgcgttacataacttacggtaaatggcccgcctggctgaccgcccaacgacccccgcccattgacg
tcaataatgacgtatgttcccatagtaacgccaatagggactttccattgacgtcaatgggtggactatttacggtaaactgcccacttggcagtacatc
aagtgtatcatatgccaagtacgccccctattgacgtcaatgacggtaaatggcccgcctggcattatgcccagtacatgaccttatgggactttcctac
ttggcagtacatctacgtattagtcatcgctattaccatgggtcgaggtgagccccacgttctgcttcactctccccatctcccccccctccccaccccca
attttgtatttatttattttttaattattttgtgcagcgatgggggcgggggggggggggcgcgcgccaggcggggcggggcggggcgaggggcggg
gcggggcgaggcggagaggtgcggcggcagccaatcagagcggcgcgctccgaaagtttccttttatggcgaggcggcggcggcggcggccctata
aaaagcgaagcgcgcggcgggcgggagtcgctgcgttgccttcgccccgtgccccgctccgcgccgcctcgcgccgcccgccccggctctgactgacc
gcgttactcccacaggtgagcgggcgggacggcccttctcctccgggctgtaattagcgcttggtttaatgacggctcgtttcttttctgtggctgcgtga
aagccttaaagggctccgggagggccctttgtgcggggggagcggctcgggggtgcgtgcgtgtgtgtgtgcgtggggagcgccgcgtgcggccc
gcgctgcccggcggctgtgagcgctgcgggcgcggcgcggggctttgtgcgctccgcgtgtgcgcgaggggagcgcggccgggggcggtgccccgcg
gtgcggggggggctgcgaggggaacaaaggctgcgtgcggggtgtgtgcgtgggggggtgagcagggggtgtgggcgcggcggtcgggctgtaacc
cccccctgcacccccctccccgagttgctgagcacggcccggcttcgggtgcggggctccgtgcggggcgtggcgcggggctcgccgtgccgggcggg
gggtggcggcaggtgggggtgccgggcggggcggggccgcctcgggccggggagggctcgggggaggggcgcggcggcccccggagcgccggcgg
ctgtcgaggcgcggcgagccgcagccattgccttttatggtaatcgtgcgagagggcgcagggacttcctttgtcccaaatctgtgcggagccgaaatc
tgggaggcgccgccgcaccccctctagcgggcgcggggcgaagcggtgcggcgccggcaggaaggaaatgggcggggagggccttcgtgcgtcgc
cgcgccgccgtccccttctccctctccagcctcggggctgtccgcggggggacggctgccttcgggggggacggggcagggcggggttcggcttctgg
cgtgtgaccggcggctctagagcctctgctaaccatgttcatgccttcttctttttcctacagctcctgggcaacgtgctggttattgtgctgtctcatcatt

ttggcaaagaattgctcgagataacttcgtatagcatacattatacgaagttatcttaagacaaacttgtttattgcagcttataatggttacaaataaa
gcaatagcatcacaaatttcacaaataaagcatttttttcactgcattctagttgtggtttgtccaaactcatcaatgtatcttaaacttgtttattgcagct
tataatggttacaaataaagcaatagcatcacaaatttcacaaataaagcatttttttcactgcattctagttgtggtttgtccaaactcatcaatgtatc
ttaaacttgtttattgcagcttataatggttacaaataaagcaatagcatcacaaatttcacaaataaagcatttttttcactgcattctagttgtggtttg
tccaaactcatcaatgtatcttagcatggacgtcataacttcgtatagcatacattatacgaagttataccggtggtaccacgatggtgagcaagggag
aggaggtcatcaaagagttcatgcgcttcaaggtgcgcatggagggctccatgaacggccacgagttcgagatcgagggcgagggcgagggccgcc
cctacgagggcacccagaccgccaagctgaaggtgaccaagggcggccccctgcccttcgcctgggacatcctgtccccccagttcatgtacggctcc
aaggcgtacgtgaagcaccccgccgacatccccgattacaagaagctgtccttccccgagggcttcaagtgggagcgcgtgatgaacttcgaggacg
gcggtctggtgaccgtgacccaggactcctccctgcaggacggcacgctgatctacaaggtgaagatgcgcggcaccaacttcccccccgacggccc
cgtaatgcagaagaagaccatgggctgggaggcctccaccgagcgcctgtaccccgcgacggcgtgctgaagggcgagatccaccaggccctgaa
gctgaaggacggcggccactacctggtggagttcaagaccatctacatggccaagaagcccgtgcaactgcccggctactactacgtggacaccaag
ctggacatcacctcccacaacgaggactacaccatcgtggaacagtacgagcgctccgagggccgccaccacctgttcctggggcatggcaccggca
gcaccggcagcggcagctccggcaccgcctcctccgaggacaacaacatggccgtcatcaaagagttcatgcgcttcaaggtgcgcatggagggctc
catgaacggccacgagttcgagatcgagggcgagggcgagggccgcccctacgagggcacccagaccgccaagctgaaggtgaccaagggcggcc
ccctgcccttcgcctgggacatcctgtccccccagttcatgtacggctccaaggcgtacgtgaagcaccccgccgacatccccgattacaagaagctgt
ccttccccgagggcttcaagtgggagcgcgtgatgaacttcgaggacggcggtctggtgaccgtgacccaggactcctccctgcaggacggcacgctg
atctacaaggtgaagatgcgcggcaccaacttcccccccgacggccccgtaatgcagaagaagaccatgggctgggaggcctccaccgagcgcctgt
accccgcgacggcgtgctgaagggcgagatccaccaggccctgaagctgaaggacggcggccgctacctggtggagttcaagaccatctacatgg
ccaagaagcccgtgcaactgcccggctactactacgtggacaccaagctggacatcacctcccacaacgaggactacaccatcgtggaacagtacga
gcgctccgagggccgccaccacctgttcctgtacggcatggacgagctgtacaagtagtaagcttcagaggttgaagatctctgtgccttctagttgcc
agccatctgttgtttgcccctcccccgtgccttccttgaccctggaaggtgccactcccactgtcctttcctaataaaatgaggaaattgcatcgcattgtc
tgagtaggtgtcattctattctggggggtggggtggggcaggacagcaagggggaggattgggaagacaatagcaggcatgctggggatgcggtgg
gctctatgggctagaagatctctgcaactggagtctttctggaagataggcgggagtcttctgggcaggcttaaaggctaacctggtgcgtggggcgtt
gtcctgcagaggaattgaacaggtgtaaaattggaggggcaagacttcccacagattttcgattgtgttgttaagtattgtaataggggcaaataagg
gaaatagactaggcactcacctggggttttatgcagcaaaactacaggttattattgcttgtgatccgccctggagaatttttcaccgaggtagattgaa
gacatgcccacccaaattttaatattcttccacttgcgatccttgctacagtatgaaattacagtatcgtgaattagaatatataagcagaattttaagca
ttttaaaagagcccagtacttcatgtctgtctctcccacttctgcagccctatcaaagggtattttagcacactcattttagtcccattttcatttgttgtact
ggcttatccaatccctagacagagcactggcattccctctctcctgatcttagaagtccgatgactcatgaaaccagacagattagtttcatacaccaca
aatcgaggctgtagctggggcctttaacattgcagttttttttattcttcagtacactttgttgattctttgccttgatcttgacttcaggttctatcaccaccc
cctcagatggtgttccacacttgggcctattcacagttcagagagctttacaacaatagatgtattgagaatccaacctaaagttcagctttttactccca
tgaatgcctctttcctt

**Left homology arm (HL):  1-796**

**Right homology arm (HR):  4930-5733**

**CAG promoter: 1061-2765**

**Floxed stop cassette: 2771-3224**

**tdTomato: 3240-4670**

bGHpA:4695-4918 Note that due the high GC conents in some regions, we used in vitro Cre excision and transfection analysis to check this donor DNA. Please see the test results in anoter file.

```
1    CCGCGTGTGA AAACACAAAT GGCGTGTTTT GGTTGGAGTG AGGCGCCTGT CAATTAACGG CTGCCGGAGT GCGCAGCCGC TGACTGCCTC GCTGTGCCCA
101  CTGGGTGGGG CGGGAGGTAG GTGGGGTGAG GCGAGCTGGA CGTGCGGGCG CGGTCGGCCT CTGGCGGGGC GGGGGAGGGG AGGGTCAGCG AAAGTGGCTG
201  GCGCGTGAGC GGCCTCCCAC CCTCCCCTTC CTCTGGGGGA GTCGTTTTAC CCGCCGCCGG CCTGGCCTCG TCATCTGATT GGCTCTCGGG GCTCAGAAAA
301  CTGGCCTTTG CAATTGGCCC GCGTTCATGC AAGTTCAGTC CCTAAGCTGG CTGGCGGGGG CGGCAGGGAG GCGCTCACAG GTTCCGGCCC TCCCCCCAGG
401  CCCCGCGCCG CAGAGTCTGG CCCCGCGCCC CTGCGCAACG TGGCAGGAAG CGCGCGCTGG GGGCGGGGAC GGGCGGTCGG TCTGAGCGGC GGGCGGGTGC
501  AAACGGGATT CCTCCTTGAG TTGTGGCACT GAGGAACGTG CTGAACAAGA CCTACATTGC ACTCCAGGGA GTGGATGAAG GAGTTGGGGC TCAGTCGGGT
601  TGTATTGGAG ACAAGAAGCA CTTGCTCTCC AAAAGTCGGT TTGAGTTATC ATTAAGGGAG CTGCAGTGGA GTAGGCGGAG AAAAGGCCGC ACCCTTCTCA
701  GGACGGGGGA GGGGAGTGTT GCAATACCTT TCTGGGAGTT CTCTGCTGCC TCCTGTCTTC TGAGGACCGC CCTGGGCCTG GAAGATTCCC TTCCCCGGAT
801  CCTTGGGGTT GCGCCTTTTC CAAGGCAGCC CTGGGTTTGC GCAGGGACGC GGCTGCTCTG GGCGTGGTTC CGGGAAACGC AGCGGCGCCG ACCCTGGGTC
901  TCGCACATTC TTCACGTCCG TTCGCAGCGT CACCCGGATC TTCGCCGCTA CCCTTGTGGG CCCCCCGGCG ACGCTTCCTG CTCCGCCCCT AAGTCGGGAA
1001 GGTTCCTTGC GGTTCGCGGC GTGCCGGACG TGACAAACGG AAGCCGCACG TCTCACTAGT TATTAATAGT AATCAATTAC GGGGTCATTA GTTCATAGCC
1101 CATATATGGA GTTCCGCGTT ACATAACTTA CGGTAAATGG CCCGCCTGGC TGACCGCCCA ACGACCCCCG CCCATTGACG TCAATAATGA CGTATGTTCC
1201 CATAGTAACG CCAATAGGGA CTTTCCATTG ACGTCAATGG GTGGACTATT TACGGTAAAC TGCCCACTTG GCAGTACATC AAGTGTATCA TATGCCAAGT
1301 ACGCCCCCTA TTGACGTCAA TGACGGTAAA TGGCCCGCCT GGCATTATGC CCAGTACATG ACCTTATGGG ACTTTCCTAC TTGGCAGTAC ATCTACGTAT
1401 TAGTCATCGC TATTACCATG GGTCGAGGTG AGCCCCACGT TCTGCTTCAC TCTCCCCATC TCCCCCCCCT CCCCACCCCC AATTTTGTAT TTATTTATTT
1501 TTTAATTATT TTGTGCAGCG ATGGGGGCGG GGGGGGGGGG GGCGCGCGCC AGGCGGGGCG GGGCGGGGCG AGGGGCGGGG CGGGGCGAGG CGGAGAGGTG
1601 CGGCGGCAGC CAATCAGAGC GGCGCGCTCC GAAAGTTTCC TTTTATGGCG AGGCGGCGGC GGCGGCGGCC CTATAAAAAG CGAAGCGCGC GGCGGGCGGG
1701 AGTCGCTGCG TTGCCTTCGC CCCGTGCCCC GCTCCGCGCC GCCTCGCGCC GCCCGCCCCG GCTCTGACTG ACCGCGTTAC TCCCACAGGT GAGCGGGCGG
1801 GACGGCCCTT CTCCTCCGGG CTGTAATTAG CGCTTGGTTT AATGACGGCT CGTTTCTTTT CTGTGGCTGC GTGAAAGCCT TAAAGGGCTC CGGGAGGGCC
1901 CTTTGTGCGG GGGGGAGCGG CTCGGGGGGT GCGTGCGTGT GTGTGTGCGT GGGGAGCGCC GCGTGCGGCC CGCGCTGCCC GGCGGCTGTG AGCGCTGCGG
2001 GCGCGGCGCG GGGCTTTGTG CGCTCCGCGT GTGCGCGAGG GGAGCGCGGC CGGGGGCGGT GCCCCGCGGT GCGGGGGGGC TGCGAGGGGA ACAAAGGCTG
2101 CGTGCGGGGT GTGTGCGTGG GGGGGTGAGC AGGGGGTGTG GGCGCGGCGG TCGGGCTGTA ACCCCCCCCT GCACCCCCCT CCCCGAGTTG CTGAGCACGG
2201 CCCGGCTTCG GGTGCGGGGC TCCGTGCGGG GCGTGGCGCG GGGCTCGCCG TGCCGGGCGG GGGGTGGCGG CAGGTGGGGG TGCCGGGCGG GGCGGGGCCG
2301 CCTCGGGCCG GGGAGGGCTC GGGGGAGGGG CGCGGCGGCC CCGGAGCGCC GGCGGCTGTC GAGGCGCGGC GAGCCGCAGC CATTGCCTTT TATGGTAATC
2401 GTGCGAGAGG GCGCAGGGAC TTCCTTTGTC CCAAATCTGT GCGGAGCCGA AATCTGGGAG GCGCCGCCGC ACCCCCTCTA GCGGGCGCGG GGCGAAGCGG
2501 TGCGGCGCCG GCAGGAAGGA AATGGGCGGG GAGGGCCTTC GTGCGTCGCC GCGCCGCCGT CCCCTTCTCC CTCTCCAGCC TCGGGGCTGT CCGCGGGGGG
2601 ACGGCTGCCT TCGGGGGGGA CGGGGCAGGG CGGGGTTCGG CTTCTGGCGT GTGACCGGCG GCTCTAGAGC CTCTGCTAAC CATGTTCATG CCTTCTTCTT
2701 TTTCCTACAG CTCCTGGGCA ACGTGCTGGT TATTGTGCTG TCTCATCATT TTGGCAAAGA ATTGCTCGAG ATAACTTCGT ATAGCATACA TTATACGAAG
2801 TTATCTTAAG ACAAACTTGT TTATTGCAGC TTATAATGGT TACAAATAAA GCAATAGCAT CACAAATTTC ACAAATAAAG CATTTTTTTC ACTGCATTCT
2901 AGTTGTGGTT TGTCCAAACT CATCAATGTA TCTTAAACTT GTTTATTGCA GCTTATAATG GTTACAAATA AAGCAATAGC ATCACAAATT TCACAAATAA
3001 AGCATTTTTT TCACTGCATT CTAGTTGTGG TTTGTCCAAA CTCATCAATG TATCTTAAAC TTGTTTATTG CAGCTTATAA TGGTTACAAA TAAAGCAATA
3101 GCATCACAAA TTTCACAAAT AAAGCATTTT TTTCACTGCA TTCTAGTTGT GGTTTGTCCA AACTCATCAA TGTATCTTAG CATGGACGTC ATAACTTCGT
+3                                                 Met Val Ser Lys Gly Glu Glu Val Ile Lys Glu Phe Met Arg Phe Lys Val Arg Met Glu Gly
3201 ATAGCATACA TTATACGAAG TTATACCGGT GGTACCACGA TGGTGAGCAA GGGAGAGGAG GTCATCAAAG AGTTCATGCG CTTCAAGGTG CGCATGGAGG
+3   Gly Ser Met Asn Gly His Glu Phe Glu Ile Glu Gly Glu Gly Glu Gly Arg Pro Tyr Glu Gly Thr Gln Thr Ala Lys Leu Lys Val Thr Lys Gly Gly Pro
3301 GCTCCATGAA CGGCCACGAG TTCGAGATCG AGGGCGAGGG CGAGGGCCGC CCCTACGAGG GCACCCAGAC CGCCAAGCTG AAGGTGACCA AGGGCGGCCC
+3   Pro Leu Pro Phe Ala Trp Asp Ile Leu Ser Pro Gln Phe Met Tyr Gly Ser Lys Ala Tyr Val Lys His Pro Ala Asp Ile Pro Asp Tyr Lys Lys Leu Ser
3401 CCTGCCCTTC GCCTGGGACA TCCTGTCCCC CCAGTTCATG TACGGCTCCA AGGCGTACGT GAAGCACCCC GCCGACATCC CCGATTACAA GAAGCTGTCC
+3   Phe Pro Glu Gly Phe Lys Trp Glu Arg Val Met Asn Phe Glu Asp Gly Gly Leu Val Thr Val Thr Gln Asp Ser Ser Leu Gln Asp Gly Thr Leu Ile Tyr
3501 TTCCCCGAGG GCTTCAAGTG GGAGCGCGTG ATGAACTTCG AGGACGGCGG TCTGGTGACC GTGACCCAGG ACTCCTCCCT GCAGGACGGC ACGCTGATCT
+3   Tyr Lys Val Lys Met Arg Gly Thr Asn Phe Pro Pro Asp Gly Pro Val Met Gln Lys Lys Thr Met Gly Trp Glu Ala Ser Thr Glu Arg Leu Tyr Pro Arg
3601 ACAAGGTGAA GATGCGCGGC ACCAACTTCC CCCCCGACGG CCCCGTAATG CAGAAGAAGA CCATGGGCTG GGAGGCCTCC ACCGAGCGCC TGTACCCCCG
+3   Arg Asp Gly Val Leu Lys Gly Glu Ile His Gln Ala Leu Lys Leu Lys Asp Gly Gly His Tyr Leu Val Glu Phe Lys Thr Ile Tyr Met Ala Lys Lys Pro
3701 CGACGGCGTG CTGAAGGGCG AGATCCACCA GGCCCTGAAG CTGAAGGACG GCGGCCACTA CCTGGTGGAG TTCAAGACCA TCTACATGGC CAAGAAGCCC
+3   Val Gln Leu Pro Gly Tyr Tyr Tyr Val Asp Thr Lys Leu Asp Ile Thr Ser His Asn Glu Asp Tyr Thr Ile Val Glu Gln Tyr Glu Arg Ser Glu Gly Arg
3801 GTGCAACTGC CCGGCTACTA CTACGTGGAC ACCAAGCTGG ACATCACCTC CCACAACGAG GACTACACCA TCGTGGAACA GTACGAGCGC TCCGAGGGCC
+3   Arg His His Leu Phe Leu Gly His Gly Thr Gly Ser Thr Gly Ser Gly Ser Ser Gly Thr Ala Ser Ser Glu Asp Asn Asn Met Ala Val Ile Lys Glu Phe
3901 GCCACCACCT GTTCCTGGGG CATGGCACCG GCAGCACCGG CAGCGGCAGC TCCGGCACCG CCTCCTCCGA GGACAACAAC ATGGCCGTCA TCAAAGAGTT
+3   Phe Met Arg Phe Lys Val Arg Met Glu Gly Ser Met Asn Gly His Glu Phe Glu Ile Glu Gly Glu Gly Glu Gly Arg Pro Tyr Glu Gly Thr Gln Thr Ala
4001 CATGCGCTTC AAGGTGCGCA TGGAGGGCTC CATGAACGGC CACGAGTTCG AGATCGAGGG CGAGGGCGAG GGCCGCCCCT ACGAGGGCAC CCAGACCGCC
+3   Lys Leu Lys Val Thr Lys Gly Gly Pro Leu Pro Phe Ala Trp Asp Ile Leu Ser Pro Gln Phe Met Tyr Gly Ser Lys Ala Tyr Val Lys His Pro Ala Asp
4101 AAGCTGAAGG TGACCAAGGG CGGCCCCCTG CCCTTCGCCT GGGACATCCT GTCCCCCCAG TTCATGTACG GCTCCAAGGC GTACGTGAAG CACCCCGCCG
+3   Asp Ile Pro Asp Tyr Lys Lys Leu Ser Phe Pro Glu Gly Phe Lys Trp Glu Arg Val Met Asn Phe Glu Asp Gly Gly Leu Val Thr Val Thr Gln Asp Ser
4201 ACATCCCCGA TTACAAGAAG CTGTCCTTCC CCGAGGGCTT CAAGTGGGAG CGCGTGATGA ACTTCGAGGA CGGCGGTCTG GTGACCGTGA CCCAGGACTC
+3   Ser Ser Leu Gln Asp Gly Thr Leu Ile Tyr Lys Val Lys Met Arg Gly Thr Asn Phe Pro Pro Asp Gly Pro Val Met Gln Lys Lys Thr Met Gly Trp Glu
4301 CTCCCTGCAG GACGGCACGC TGATCTACAA GGTGAAGATG CGCGGCACCA ACTTCCCCCC CGACGGCCCC GTAATGCAGA AGAAGACCAT GGGCTGGGAG
+3   Ala Ser Thr Glu Arg Leu Tyr Pro Arg Asp Gly Val Leu Lys Gly Glu Ile His Gln Ala Leu Lys Leu Lys Asp Gly Gly Arg Tyr Leu Val Glu Phe Lys
4401 GCCTCCACCG AGCGCCTGTA CCCCCGCGAC GGCGTGCTGA AGGGCGAGAT CCACCAGGCC CTGAAGCTGA AGGACGGCGG CCGCTACCTG GTGGAGTTCA
+3   Lys Thr Ile Tyr Met Ala Lys Lys Pro Val Gln Leu Pro Gly Tyr Tyr Tyr Val Asp Thr Lys Leu Asp Ile Thr Ser His Asn Glu Asp Tyr Thr Ile Val
4501 AGACCATCTA CATGGCCAAG AAGCCCGTGC AACTGCCCGG CTACTACTAC GTGGACACCA AGCTGGACAT CACCTCCCAC AACGAGGACT ACACCATCGT
+3   Val Glu Gln Tyr Glu Arg Ser Glu Gly Arg His His Leu Phe Leu Tyr Gly Met Asp Glu Leu Tyr Lys ***
4601 GGAACAGTAC GAGCGCTCCG AGGGCCGCCA CCACCTGTTC CTGTACGGCA TGGACGAGCT GTACAAGTAG TAAGCTTCAG AGGTTGAAGA TCTCTGTGCC
4701 TTCTAGTTGC CAGCCATCTG TTGTTTGCCC CTCCCCCGTG CCTTCCTTGA CCCTGGAAGG TGCCACTCCC ACTGTCCTTT CCTAATAAAA TGAGGAAATT
4801 GCATCGCATT GTCTGAGTAG GTGTCATTCT ATTCTGGGGG GTGGGGTGGG GCAGGACAGC AAGGGGGAGG ATTGGGAAGA CAATAGCAGG CATGCTGGGG
4901 ATGCGGTGGG CTCTATGGGC TAGAAGATCT CTGCAACTGG AGTCTTTCTG GAAGATAGGC GGGAGTCTTC TGGGCAGGCT TAAAGGCTAA CCTGGTGCGT
5001 GGGGCGTTGT CCTGCAGAGG AATTGAACAG GTGTAAAATT GGAGGGGCAA GACTTCCCAC AGATTTTCGA TTGTGTTGTT AAGTATTGTA ATAGGGGCAA
5101 ATAAGGGAAA TAGACTAGGC ACTCACCTGG GGTTTTATGC AGCAAAACTA CAGGTTATTA TTGCTTGTGA TCCGCCCTGG AGAATTTTTC ACCGAGGTAG
5201 ATTGAAGACA TGCCCACCCA AATTTTAATA TTCTTCCACT TGCGATCCTT GCTACAGTAT GAAATTACAG TATCGTGAAT TAGAATATAT AAGCAGAATT
5301 TTAAGCATTT TAAAAGAGCC CAGTACTTCA TGTCTGTCTC TCCCACTTCT GCAGCCCTAT CAAAGGGTAT TTTAGCACAC TCATTTTAGT CCCATTTTCA
5401 TTTGTTGTAC TGGCTTATCC AATCCCTAGA CAGAGCACTG GCATTCCCTC TCTCCTGATC TTAGAAGTCC GATGACTCAT GAAACCAGAC AGATTAGTTT
5501 CATACACCAC AAATCGAGGC TGTAGCTGGG GCCTTTAACA TTGCAGTTTT TTTATTCTTC AGTACACTTT GTTGATTCTT TGCCTTGATC TTGACTTCAG
5601 GTTCTATCAC CACCCCCTCA GATGGTGTTC CACACTTGGG CCTATTCACA GTTCAGAGAG CTTTACAACA ATAGATGTAT TGAGAATCCA ACCTAAAGTT
5701 CAGCTTTTTA CTCCCATGAA TGCCTCTTTC CTT
```


Designed by: **Guojun (Justin) Zhao, Ph.D.**

Envigo+++

2033 Westport Center Drive, Saint Louis, MO 63146, USA
