## Supplementary material for "Generation of a new immunodeficient rat model of retinal degeneration with LSL TdTomato reporter and TdTomato-Pcp2 expression": 2. Donor Design_ Pcp2-1105 donor DNA.pdf

**Donor DNA Design**

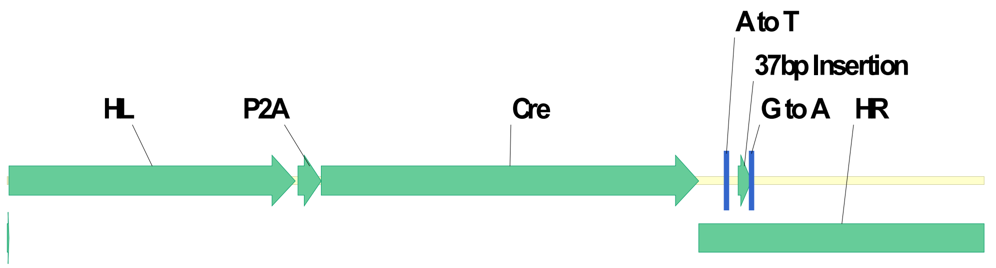


Pcp2-1105 Donor DNA Design

2731bp

**Sequence (2731bp):**

TTTGGGGGCGTTCCACTTTATTGTTTCTGAGACAGGGTTTCATGGAGCTCTGGCTGGCCAACGATGACCTTAAACT
TCTACTTTTCCAACATGTACATCCTGGGTTCTGGGATTACAGGTGCGAGCCACTGTGAGTGATTTATGTGGTTAGG
GATGGAATCCGGGACTTTGATGCAGTCCCTATTAACTGAATCCCGGCCCTGGCCCTATGTCTGTGTACACTGGCAG
ACACGTTCAGAGGTCCATGTGCAGATGGGTACACGTGCAGGTAGAAGCCAGAGGTTCTTCGGGAGTCATCACCTT
GTTATTTGAGATGGGATCCTCCACTGCCCTGGAACTCACCAGGCAGATCAGGTTCCCTAGCTAGGAAGCCCTAGG
GACCCTCCTGACTCTCCGTCCTAAGCACTGGGATTACAAGTGCCTGAAGCCACACCTGGCCTCCCAGGATCCAAAT
GAGATCTTCAAACGAGCACAGCAAGAATTGCAATGACTGAGCTATATTCTCAGCGCCCAACATCTGTTTTAGTATG
TAGACCAGGGTGGCTTCAAACTCGAGAAAATCCTCCTGCCTCTGTTTCCTGAGTACTCACGACCACCCTGATTGGC
AGACGTAAAGACATGTTGTTTGGAGGTGCCCCTGACTTGCACTTTCCGTGGTCCCCTCCCACCGTCTGACCCTTCTT
CATCCACAGGATGGAATGCAGAAACGACCCGGGACACTCAGCCCTCAACCCCTGCTCAGCCCGCAGGACCCTGCT
GCTCTCAGCTTCCGCAGGAACAGTAGCCCCCAGCCCCAGACACAGGCTCCAGGAAGCGGAGCTACTAACTTCAGC
CTGCTGAAGCAGGCTGGAGACGTGGAGGAGAACCCTGGACCTATGGCACCCAAGAAGAAGAGGAAGGTGTCCA
ATTTACTGACCGTACACCAAAATTTGCCTGCATTACCGGTCGATGCAACGAGTGATGAGGTTCGCAAGAACCTGAT
GGACATGTTCAGGGATCGCCAGGCGTTTTCTGAGCATACCTGGAAAATGCTTCTGTCCGTTTGCCGGTCGTGGGC
GGCATGGTGCAAGTTGAATAACCGGAAATGGTTTCCCGCAGAACCTGAAGATGTTCGCGATTATCTTCTATATCTT
CAGGCGCGCGGTCTGGCAGTAAAAACTATCCAGCAACATTTGGGCCAGCTAAACATGCTTCATCGTCGGTCCGGG
CTGCCACGACCAAGTGACAGCAATGCTGTTTCACTGGTTATGCGGCGGATCCGAAAAGAAAACGTTGATGCCGGT
GAACGTGCAAAACAGGCTCTAGCGTTCGAACGCACTGATTTCGACCAGGTTCGTTCACTCATGGAAAATAGCGATC
GCTGCCAGGATATACGTAATCTGGCATTTCTGGGGATTGCTTATAACACCCTGTTACGTATAGCCGAAATTGCCAG
GATCAGGGTTAAAGATATCTCACGTACTGACGGTGGGAGAATGTTAATCCATATTGGCAGAACGAAAACGCTGGT
TAGCACCGCAGGTGTAGAGAAGGCACTTAGCCTGGGGGTAACTAAACTGGTCGAGCGATGGATTTCCGTCTCTGG
TGTAGCTGATGATCCGAATAACTACCTGTTTTGCCGGGTCAGAAAAAATGGTGTTGCCGCGCCATCTGCCACCAGC
CAGCTATCAACTCGCGCCCTGGAAGGGATTTTTGAAGCAACTCATCGATTGATTTACGGCGCTAAGGATGACTCTG

GTCAGAGATACCTGGCCTGGTCTGGACACAGTGCCCGTGTCGGAGCCGCGCGAGATATGGCCCGCGCTGGAGTTT
CAATACCGGAGATCATGCAAGCTGGTGGCTGGACCAATGTAAATATTGTCATGAACTATATCCGTAACCTGGATAG
TGAAACAGGGGCAATGGTGCGCCTGCTGGAAGATGGCGATTGACACAAGCTCCCTGAGAGTTCCAGCCACCCTG
GGCCTCCCACTGGCCCCTGAAAGTAATAAAATACTTGGCACTAGCGACTGAGAGTTCAGTGTGTGTTATTTCCTGG
GGGCAGGGTCTGAGAGTTCAGTGTGTGTTATTTCCTGGGGAGGGGGGTGTCTGGAACTTGAGGAACTGAAGGTC
TCAGGAGCTCTGCTGGGCAGCTTGAAGAAGTCTTTCTTCCCTCTGCTTCCGGATCTTCTGTTTAAATTCCTCTAGCTC
CTGGCGCTGGAATGGGGAGAGGGGTGTGATGGAAAGGAAGGAGGGGTATAGGCCTCTGGGCCTGGACTTACTC
ACGTTACCCTCCCTTTGGCTTTAGAGTCCAGTATTCACACTAGGGACCCCACGCCAATCACAACCACCATACACACG
TGAGTTCAGCTTTATCCCTCAGGGAAAAGGTCATTTACGAAACCATTTGTGCCCTCCCCCTCTTTTTAAGATGGGGT
CTCATATACTACAGGCTAGCCTTGAGCTCACTGTGTGGCACAGAACAGCCAGAATCCTCCCCATCCTCCTGCATCCA
TCTCCTGAGTGCTAGAATGCTGGATTTACGGCTACCTCTCCTGTGATCATGGGTATATACCATCACAAACGGCAAG
AAACTTTTACGGAGACAAGGTCCCAGGCTAGTCTGGAACTCCTACTCCTTCCACTTCCACTTCCCTATCTAGGGCTG
AGGGTGAAATCTTCATTTGGGATAAAGTGACCCAGTCCCTTAGGGAGAACAGGTTACTGACCTGGCCCTCTTTCTC
TCGTGGCC

**Homology arm-left(HL): 1-802**

**P2A: 810-875**

**Cre cDNA: 876-1931**

**Homology arm-right (HR): 1931-2731**

```
1     TTTGGGGGCG TTCCACTTTA TTGTTTCTGA GACAGGGTTT CATGGAGCTC TGGCTGGCCA ACGATGACCT TAAACTTCTA CTTTTCCAAC ATGTACATCC
      AAACCCCCGC AAGGTGAAAT AACAAAGACT CTGTCCCAAA GTACCTCGAG ACCGACCGGT TGCTACTGGA ATTTGAAGAT GAAAAGGTTG TACATGTAGG
101   TGGGTTCTGG GATTACAGGT GCGAGCCACT GTGAGTGATT TATGTGGTTA GGGATGGAAT CCGGGACTTT GATGCAGTCC CTATTAACTG AATCCCGGCC
      ACCCAAGACC CTAATGTCCA CGCTCGGTGA CACTCACTAA ATACACCAAT CCCTACCTTA GGCCCTGAAA CTACGTCAGG GATAATTGAC TTAGGGCCGG
201   CTGGCCCTAT GTCTGTGTAC ACTGGCAGAC ACGTTCAGAG GTCCATGTGC AGATGGGTAC ACGTGCAGGT AGAAGCCAGA GGTTCTTCGG GAGTCATCAC
      GACCGGGATA CAGACACATG TGACCGTCTG TGCAAGTCTC CAGGTACACG TCTACCCATG TGCACGTCCA TCTTCGGTCT CCAAGAAGCC CTCAGTAGTG
301   CTTGTTATTT GAGATGGGAT CCTCCACTGC CCTGGAACTC ACCAGGCAGA TCAGGTTCCC TAGCTAGGAA GCCCTAGGGA CCCTCCTGAC TCTCCGTCCT
      GAACAATAAA CTCTACCCTA GGAGGTGACG GGACCTTGAG TGGTCCGTCT AGTCCAAGGG ATCGATCCTT CGGGATCCCT GGGAGGACTG AGAGGCAGGA
401   AAGCACTGGG ATTACAAGTG CCTGAAGCCA CACCTGGCCT CCCAGGATCC AAATGAGATC TTCAAACGAG CACAGCAAGA ATTGCAATGA CTGAGCTATA
      TTCGTGACCC TAATGTTCAC GGACTTCGGT GTGGACCGGA GGGTCCTAGG TTTACTCTAG AAGTTTGCTC GTGTCGTTCT TAACGTTACT GACTCGATAT
501   TTCTCAGCGC CCAACATCTG TTTTAGTATG TAGACCAGGG TGGCTTCAAA CTCGAGAAAA TCCTCCTGCC TCTGTTTCCT GAGTACTCAC GACCACCCTG
      AAGAGTCGCG GGTTGTAGAC AAAATCATAC ATCTGGTCCC ACCGAAGTTT GAGCTCTTTT AGGAGGACGG AGACAAAGGA CTCATGAGTG CTGGTGGGAC
+3                                                                                                     Asp  Gly  Met
601   ATTGGCAGAC GTAAAGACAT GTTGTTTGGA GGTGCCCCTG ACTTGCACTT TCCGTGGTCC CCTCCCACCG TCTGACCCTT CTTCATCCAC AGGATGGAAT
      TAACCGTCTG CATTTCTGTA CAACAAACCT CCACGGGGAC TGAACGTGAA AGGCACCAGG GGAGGGTGGC AGACTGGGAA GAAGTAGGTG TCCTACCTTA
+3    Met Gln Lys Arg Pro Gly Thr Leu Ser Pro Gln Pro Leu Leu Ser Pro Gln Asp Pro Ala Ala Leu Ser Phe Arg Arg Asn Ser Ser Pro Gln Pro Gln Thr
701   GCAGAAACGA CCCGGGACAC TCAGCCCTCA ACCCCTGCTC AGCCCGCAGG ACCCTGCTGC TCTCAGCTTC CGCAGGAACA GTAGCCCCCA GCCCCAGACA
      CGTCTTTGCT GGGCCCTGTG AGTCGGGAGT TGGGGACGAG TCGGGCGTCC TGGGACGACG AGAGTCGAAG GCGTCCTTGT CATCGGGGGT CGGGGTCTGT
+3    Gln Ala Pro Gly Ser Gly Ala Thr Asn Phe Ser Leu Leu Lys Gln Ala Gly Asp Val Glu Glu Asn Pro Gly Pro Met Ala Pro Lys Lys Lys Arg Lys Val
801   CAGGCTCCAG GAAGCGGAGC TACTAACTTC AGCCTGCTGA AGCAGGCTGG AGACGTGGAG GAGAACCCTG GACCTATGGC ACCCAAGAAG AAGAGGAAGG
      GTCCGAGGTC CTTCGCCTCG ATGATTGAAG TCGGACGACT TCGTCCGACC TCTGCACCTC CTCTTGGGAC CTGGATACCG TGGGTTCTTC TTCTCCTTCC
+3    Val Ser Asn Leu Leu Thr Val His Gln Asn Leu Pro Ala Leu Pro Val Asp Ala Thr Ser Asp Glu Val Arg Lys Asn Leu Met Asp Met Phe Arg Asp Arg
901   TGTCCAATTT ACTGACCGTA CACCAAAATT TGCCTGCATT ACCGGTCGAT GCAACGAGTG ATGAGGTTCG CAAGAACCTG ATGGACATGT TCAGGGATCG
      ACAGGTTAAA TGACTGGCAT GTGGTTTTAA ACGGACGTAA TGGCCAGCTA CGTTGCTCAC TACTCCAAGC GTTCTTGGAC TACCTGTACA AGTCCCTAGC
+3    Arg Gln Ala Phe Ser Glu His Thr Trp Lys Met Leu Leu Ser Val Cys Arg Ser Trp Ala Ala Trp Cys Lys Leu Asn Asn Arg Lys Trp Phe Pro Ala Glu
1001  CCAGGCGTTT TCTGAGCATA CCTGGAAAAT GCTTCTGTCC GTTTGCCGGT CGTGGGCGGC ATGGTGCAAG TTGAATAACC GGAAATGGTT TCCCGCAGAA
      GGTCCGCAAA AGACTCGTAT GGACCTTTTA CGAAGACAGG CAAACGGCCA GCACCCGCCG TACCACGTTC AACTTATTGG CCTTTACCAA AGGGCGTCTT
+3    Pro Glu Asp Val Arg Asp Tyr Leu Leu Tyr Leu Gln Ala Arg Gly Leu Ala Val Lys Thr Ile Gln Gln His Leu Gly Gln Leu Asn Met Leu His Arg Arg
1101  CCTGAAGATG TTCGCGATTA TCTTCTATAT CTTCAGGCGC GCGGTCTGGC AGTAAAAACT ATCCAGCAAC ATTTGGGCCA GCTAAACATG CTTCATCGTC
      GGACTTCTAC AAGCGCTAAT AGAAGATATA GAAGTCCGCG CGCCAGACCG TCATTTTTGA TAGGTCGTTG TAAACCCGGT CGATTTGTAC GAAGTAGCAG
+3    Arg Ser Gly Leu Pro Arg Pro Ser Asp Ser Asn Ala Val Ser Leu Val Met Arg Arg Ile Arg Lys Glu Asn Val Asp Ala Gly Glu Arg Ala Lys Gln Ala
1201  GGTCCGGGCT GCCACGACCA AGTGACAGCA ATGCTGTTTC ACTGGTTATG CGGCGGATCC GAAAAGAAAA CGTTGATGCC GGTGAACGTG CAAAACAGGC
      CCAGGCCCGA CGGTGCTGGT TCACTGTCGT TACGACAAAG TGACCAATAC GCCGCCTAGG CTTTTCTTTT GCAACTACGG CCACTTGCAC GTTTTGTCCG
+3    Ala Leu Ala Phe Glu Arg Thr Asp Phe Asp Gln Val Arg Ser Leu Met Glu Asn Ser Asp Arg Cys Gln Asp Ile Arg Asn Leu Ala Phe Leu Gly Ile Ala
1301  TCTAGCGTTC GAACGCACTG ATTTCGACCA GGTTCGTTCA CTCATGGAAA ATAGCGATCG CTGCCAGGAT ATACGTAATC TGGCATTTCT GGGGATTGCT
      AGATCGCAAG CTTGCGTGAC TAAAGCTGGT CCAAGCAAGT GAGTACCTTT TATCGCTAGC GACGGTCCTA TATGCATTAG ACCGTAAAGA CCCCTAACGA
+3    Tyr Asn Thr Leu Leu Arg Ile Ala Glu Ile Ala Arg Ile Arg Val Lys Asp Ile Ser Arg Thr Asp Gly Gly Arg Met Leu Ile His Ile Gly Arg Thr Lys
1401  TATAACACCC TGTTACGTAT AGCCGAAATT GCCAGGATCA GGGTTAAAGA TATCTCACGT ACTGACGGTG GGAGAATGTT AATCCATATT GGCAGAACGA
      ATATTGTGGG ACAATGCATA TCGGCTTTAA CGGTCCTAGT CCCAATTTCT ATAGAGTGCA TGACTGCCAC CCTCTTACAA TTAGGTATAA CCGTCTTGCT
+3    Lys Thr Leu Val Ser Thr Ala Gly Val Glu Lys Ala Leu Ser Leu Gly Val Thr Lys Leu Val Glu Arg Trp Ile Ser Val Ser Gly Val Ala Asp Asp Pro
1501  AAACGCTGGT TAGCACCGCA GGTGTAGAGA AGGCACTTAG CCTGGGGGTA ACTAAACTGG TCGAGCGATG GATTTCCGTC TCTGGTGTAG CTGATGATCC
      TTTGCGACCA ATCGTGGCGT CCACATCTCT TCCGTGAATC GGACCCCCAT TGATTTGACC AGCTCGCTAC CTAAAGGCAG AGACCACATC GACTACTAGG
+3    Pro Asn Asn Tyr Leu Phe Cys Arg Val Arg Lys Asn Gly Val Ala Ala Pro Ser Ala Thr Ser Gln Leu Ser Thr Arg Ala Leu Glu Gly Ile Phe Glu Ala
1601  GAATAACTAC CTGTTTTGCC GGGTCAGAAA AAATGGTGTT GCCGCGCCAT CTGCCACCAG CCAGCTATCA ACTCGCGCCC TGGAAGGGAT TTTTGAAGCA
      CTTATTGATG GACAAAACGG CCCAGTCTTT TTTACCACAA CGGCGCGGTA GACGGTGGTC GGTCGATAGT TGAGCGCGGG ACCTTCCCTA AAAACTTCGT
+3    Thr His Arg Leu Ile Tyr Gly Ala Lys Asp Asp Ser Gly Gln Arg Tyr Leu Ala Trp Ser Gly His Ser Ala Arg Val Gly Ala Ala Arg Asp Met Ala Arg
1701  ACTCATCGAT TGATTTACGG CGCTAAGGAT GACTCTGGTC AGAGATACCT GGCCTGGTCT GGACACAGTG CCCGTGTCGG AGCCGCGCGA GATATGGCCC
      TGAGTAGCTA ACTAAATGCC GCGATTCCTA CTGAGACCAG TCTCTATGGA CCGGACCAGA CCTGTGTCAC GGGCACAGCC TCGGCGCGCT CTATACCGGG
+3    Arg Ala Gly Val Ser Ile Pro Glu Ile Met Gln Ala Gly Gly Trp Thr Asn Val Asn Ile Val Met Asn Tyr Ile Arg Asn Leu Asp Ser Glu Thr Gly Ala
1801  GCGCTGGAGT TTCAATACCG GAGATCATGC AAGCTGGTGG CTGGACCAAT GTAAATATTG TCATGAACTA TATCCGTAAC CTGGATAGTG AAACAGGGGC
      CGCGACCTCA AAGTTATGGC CTCTAGTACG TTCGACCACC GACCTGGTTA CATTTATAAC AGTACTTGAT ATAGGCATTG GACCTATCAC TTTGTCCCCG
+3    Ala Met Val Arg Leu Leu Glu Asp Gly Asp ***
1901  AATGGTGCGC CTGCTGGAAG ATGGCGATTG ACACAAGCTC CCTGAGAGTT CCAGCCACCC TGGGCCTCCC ACTGGCCCCT GAAAGTAATA AAATACTTGG
      TTACCACGCG GACGACCTTC TACCGCTAAC TGTGTTCGAG GGACTCTCAA GGTCGGTGGG ACCCGGAGGG TGACCGGGGA CTTTCATTAT TTTATGAACC
2001  CACTAGCGAC TGAGAGTTCA GTGTGTGTTA TTTCCTGGGG GCAGGGTCTG AGAGTTCAGT GTGTGTTATT TCCTGGGGAG GGGGGTGTCT GGAACTTGAG
      GTGATCGCTG ACTCTCAAGT CACACACAAT AAAGGACCCC CGTCCCAGAC TCTCAAGTCA CACACAATAA AGGACCCCTC CCCCCACAGA CCTTGAACTC
2101  GAACTGAAGG TCTCAGGAGC TCTGCTGGGC AGCTTGAAGA AGTCTTTCTT CCCTCTGCTT CCGGATCTTC TGTTTAAATT CCTCTAGCTC CTGGCGCTGG
      CTTGACTTCC AGAGTCCTCG AGACGACCCG TCGAACTTCT TCAGAAAGAA GGGAGACGAA GGCCTAGAAG ACAAATTTAA GGAGATCGAG GACCGCGACC
2201  AATGGGGAGA GGGGTGTGAT GGAAAGGAAG GAGGGGTATA GGCCTCTGGG CCTGGACTTA CTCACGTTAC CCTCCCTTTG GCTTTAGAGT CCAGTATTCA
      TTACCCCTCT CCCCACACTA CCTTTCCTTC CTCCCCATAT CCGGAGACCC GGACCTGAAT GAGTGCAATG GGAGGGAAAC CGAAATCTCA GGTCATAAGT
2301  CACTAGGGAC CCCACGCCAA TCACAACCAC CATACACACG TGAGTTCAGC TTTATCCCTC AGGGAAAAGG TCATTTACGA AACCATTTGT GCCCTCCCCC
      GTGATCCCTG GGGTGCGGTT AGTGTTGGTG GTATGTGTGC ACTCAAGTCG AAATAGGGAG TCCCTTTTCC AGTAAATGCT TTGGTAAACA CGGGAGGGGG
2401  TCTTTTTAAG ATGGGGTCTC ATATACTACA GGCTAGCCTT GAGCTCACTG TGTGGCACAG AACAGCCAGA ATCCTCCCCA TCCTCCTGCA TCCATCTCCT
      AGAAAAATTC TACCCCAGAG TATATGATGT CCGATCGGAA CTCGAGTGAC ACACCGTGTC TTGTCGGTCT TAGGAGGGGT AGGAGGACGT AGGTAGAGGA
2501  GAGTGCTAGA ATGCTGGATT TACGGCTACC TCTCCTGTGA TCATGGGTAT ATACCATCAC AAACGGCAAG AAACTTTTAC GGAGACAAGG TCCCAGGCTA
      CTCACGATCT TACGACCTAA ATGCCGATGG AGAGGACACT AGTACCCATA TATGGTAGTG TTTGCCGTTC TTTGAAAATG CCTCTGTTCC AGGGTCCGAT
2601  GTCTGGAACT CCTACTCCTT CCACTTCCAC TTCCCTATCT AGGGCTGAGG GTGAAATCTT CATTTGGGAT AAAGTGACCC AGTCCCTTAG GGAGAACAGG
      CAGACCTTGA GGATGAGGAA GGTGAAGGTG AAGGGATAGA TCCCGACTCC CACTTTAGAA GTAAACCCTA TTTCACTGGG TCAGGGAATC CCTCTTGTCC
2701  TTACTGACCT GGCCCTCTTT CTCTCGTGGC C
      AATGACTGGA CCGGGAGAAA GAGAGCACCG G
```


Designed by: **Guojun (Justin) Zhao, Ph.D.**

Envigo+++

2033 Westport Center Drive, Saint Louis, MO 63146, USA

**Expected Chimeric MUC13 Protein Seqeunce (570aa)**

GSGATNFSLLKQAGDVEENPGPMAPKKKRKVSNLLTVHQNLPALPVDATSDEVRKNLMDMFRDRQAFSEHTWKMLL
SVCRSWAAWCKLNNRKWFPAEPEDVRDYLLYLQARGLAVKTIQQHLGQLNMLHRRSGLPRPSDSNAVSLVMRRIRKE
NVDAGERAKQALAFERTDFDQVRSLMENSDRCQDIRNLAFLGIAYNTLLRIAEIARIRVKDISRTDGGRMLIHIGRTKTLV
STAGVEKALSLGVTKLVERWISVSGVADDPNNYLFCRVRKNGVAAPSATSQLSTRALEGIFEATHRLIYGAKDDSGQRYL
AWSGHSARVGAARDMARAGVSIPEIMQAGGWTNVNIVMNYIRNLDSETGAMVRLLEDGD*
